# A Systematic Evaluation of Single-Cell Batch Integration Metrics and sBEE, A New Metric

**DOI:** 10.64898/2026.04.22.720135

**Authors:** Mekan Myradov, Aissa Houdjedj, Oznur Tastan, Hilal Kazan

**Affiliations:** Faculty of Engineering and Natural Sciences, Sabanci University, Universite Caddesi, 34956, Istanbul, Turkey.; Department of Computer Engineering, Akdeniz University, Dumlupınar Bulvarı, 07058, Antalya, Turkey.; Department of Computer Engineering, Antalya Bilim University, Universite Caddesi, 07190, Antalya, Turkey.

**Keywords:** single-cell RNA sequencing, batch effect correction, batch integration, benchmarking, evaluation metric

## Abstract

**Background:** Single-cell RNA sequencing (scRNA-seq) datasets generated across laboratories and experimental conditions often exhibit batch effects that obscure biological variation. Numerous computational methods have been developed to integrate such datasets, making robust benchmarking essential. Evaluation metrics play a central role in assessing integration quality. However, existing metrics each capture only specific aspects of batch mixing and often rely on assumptions that are violated in practice, such as every cell type being present in every batch, balanced batch composition, or simple cluster geometries. Consequently, benchmarking studies frequently report discordant rankings of integration methods, complicating interpretation and method selection.

**Results:** We systematically evaluate widely used batch integration metrics using controlled synthetic scenarios that isolate common integration challenges, including imbalanced batch composition, partial cell-type overlap, heterogeneous cluster densities, and varying cluster geometries. Our analysis reveals distinct strengths, limitations, and failure modes, demonstrating that assessments of batch mixing can change substantially across different data characteristics. These observations motivated the development of sBEE (single-cell Batch Effect Evaluator), a metric that assesses batch mixing at the single-cell level using two complementary measures: local neighborhood composition and cross-batch distance relationships. Across diverse simulated scenarios, sBEE produces stable, interpretable evaluations and remains robust to the failure modes identified in existing metrics. We further validate these findings on multiple real-world scRNA-seq datasets.

**Conclusions:** Our work provides a comprehensive characterization of the behavior of widely used batch integration metrics and introduces sBEE, a robust metric that combines complementary neighborhood and distance-based information to provide more reliable assessments of batch mixing. By identifying the conditions under which existing metrics succeed or fail and providing a unified alternative, sBEE enables more consistent benchmarking of single-cell integration methods. Code and datasets are available at https://github.com/tastanlab/sBEE.

## 1 Background

As single-cell RNA sequencing (scRNA-seq) becomes widely applicable, the need for combining data generated across laboratories, platforms, and cohorts has grown [1]. Consequently, batch effects—systematic technical differences between datasets—have become a major challenge [2, 3]. When left uncorrected, batch effects can confound meaningful biological variation and lead to misleading downstream analyses [4, 5]. To mitigate batch effects, a wide range of computational methods has been developed to mitigate batch effects, including MNN [6], Harmony [7], Seurat v3 [8], scVI [9], SCI-TUNA [10], and others [11–14]. Evaluating the performance of these batch integration methods is critical [1, 10, 15]. The evaluation metrics used to assess integration quality are central to these benchmarking efforts.

Several metrics are widely used to assess batch integration quality. Luecken et al. [15] categorize batch integration metrics into two complementary classes: biological conservation and batch correction. Biological conservation metrics evaluate whether meaningful biological structure, such as cell-type identity or trajectory organization, is preserved after integration [7, 16–18]. On the other hand, batch correction metrics assess the extent to which cells from different batches are mixed in the integrated representation [7, 19]. While biological conservation can often be evaluated using known labels, batch mixing is harder to quantify because the correct degree of mixing must be inferred from how cells from different batches are distributed in the integrated space. There are a number of metrics for evaluating batch correction performance. Distance-based metrics such as the average silhouette width (ASW) [15, 19, 20] quantify the geometric separation of batches within biological groups. Neighborhood-based metrics, including the local inverse Simpson index (iLISI) [7] and the *k*-nearest neighbor batch effect test (kBET) [19], evaluate the extent of batch mixing within local cell neighborhoods. Other approaches assess complementary aspects of integration quality, such as graph connectivity (GC) [15], which measures connectivity among cells of the same type in graph-based representations, and principal component regression (PCR) [19], which quantifies residual batch-associated variation in low-dimensional embeddings. These metrics form the foundation of current benchmarking frameworks [15, 21] and are frequently used to guide method selection in large-scale atlas projects [22]. More recently, refinements of existing metrics have been proposed, including the batch-removal-adapted silhouette (BRAS) and the cell-type integration local inverse Simpson index (CiLISI) [23, 24]. These newer metrics have not yet been systematically evaluated alongside the established benchmarking metrics; here, we include them in our comparative analysis.

Although these metrics are routinely used to evaluate integration performance, several studies have reported that their behavior can depend on dataset properties, integration scenarios, and parameter choices [7, 19, 24–27]. Lengyel and Botta-Dukát [26] show that silhouette-based metrics implicitly favor compact, isotropic clusters and become unreliable when clusters are non-convex or differ substantially in scale; Rautenstrauch and Ohler [24] further demonstrate that their batch-mixing variants fail under nested batch effects, where considering only the nearest neighboring batch can yield near-maximal scores even when integration is unsuccessful. Neighborhood-based mixing scores are sensitive to their parameterization: Korsunsky et al. [7] note that the inverse Simpson’s index depends on the chosen perplexity and degrades when batches differ greatly in size, and Mandric et al. [27] show that its values, along with those of related neighbor-based scores, shift with the number of principal components retained and the choice of neighborhood size. Finally, Büttner et al. [19] tests whether neighborhood-level batch proportions match the global distribution—an assumption that is violated when cell-type composition is imbalanced across batches and can lead to spurious rejection. Such observations, however, remain fragmented across individual metrics, datasets, and scenarios. As a result, the behavior of these metrics has not been characterized systematically across the conditions commonly encountered in practice, which hinders the comparison of integration methods.

To systematically characterize these behaviors, we construct controlled evaluation scenarios that isolate key integration challenges. These include imbalanced batch composition, rare cell populations containing fewer cells than the neighborhood size used by local metrics, partial cell-type overlap across batches, and varying cluster geometries. By analyzing metric responses across these scenarios, we identify consistent failure modes that are not apparent from aggregate benchmark scores alone. This analysis provides a principled framework for understanding when existing metrics provide reliable guidance and why they frequently yield conflicting assessments of the same integration result. Guided by these insights, we also propose the single-cell Batch Effect Evaluator (sBEE), a metric that jointly incorporates local neighborhood composition and distance structure to assess batch integration quality.

## 2 Results

We evaluated sBEE and commonly used batch-mixing metrics using controlled simulations and real-world scRNA-seq benchmarks. The simulations isolate specific properties of batch structure, while the real-world benchmarks assess whether the observed metric behaviors persist in practical integration settings. Together, these experiments reveal several systematic failure modes of existing metrics. For each simulated scenario, we report scores for each cell type together with their macro-average (unweighted mean), computed as the unweighted mean of the cell-type-specific scores.

### 2.1 Evaluations under controlled simulated scenarios

We first considered a near-perfect mixing scenario with two cell types and three batches. Cell type 1 is absent from batch 3, whereas cell type 2 is present in all three batches (Scenario A, Fig. 2). sBEE, ASW, BRAS, kBET, and GC all yield scores close to 1.00, correctly indicating the absence of batch effects. In contrast, iLISI assigns a low score to cell type 1 (0.43) because this cell type is present in only two batches, making it impossible for batch 3 to appear in its local neighborhoods. Even for cell type 2, iLISI reaches only 0.81. CiLISI and PCR similarly yield scores of only 0.66 and 0.85, respectively, underestimating the integration quality.

**Fig. 1:**
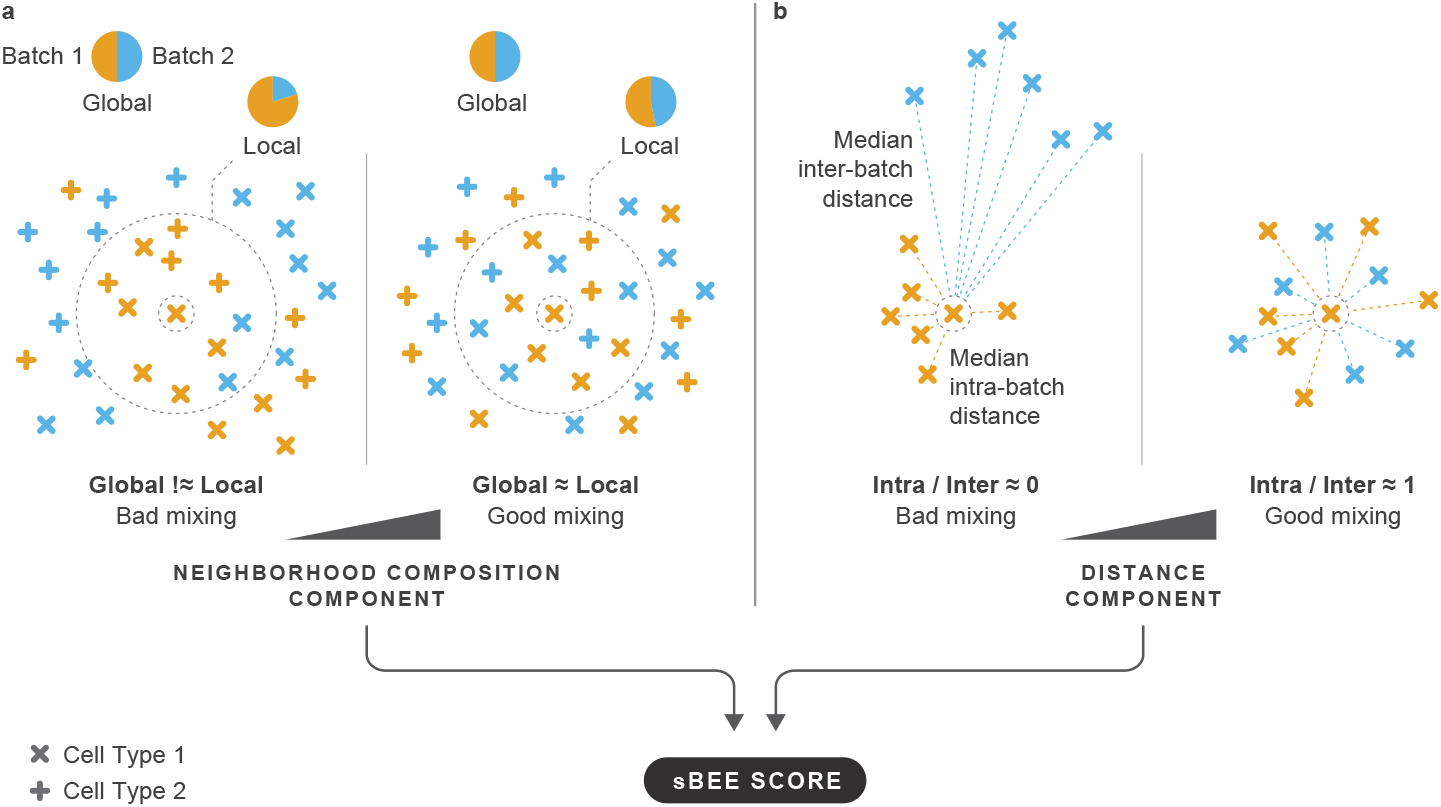
Overview of the sBEE metric computation.

**Fig. 2:**
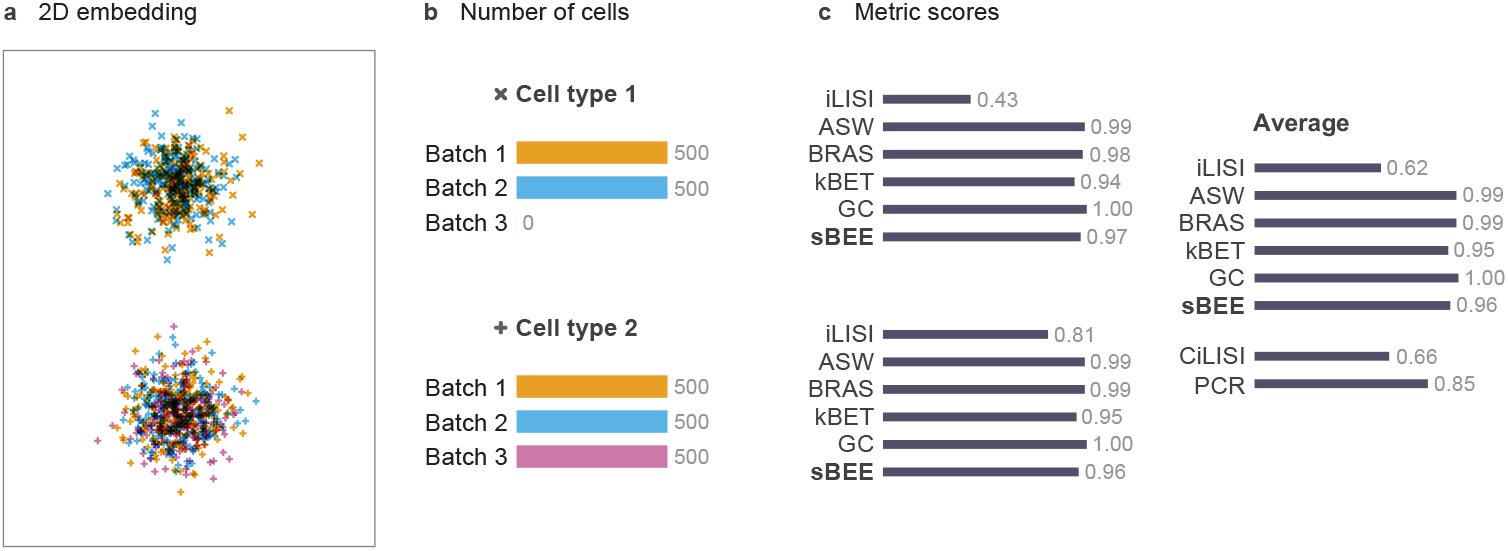
a) Simulated 2D embedding for Scenario A (near-perfect mixing). b) Batch composition of each cell type, where colors represent batches. c) Metric scores. Each cell type has a separate subplot showing its iLISI, ASW, BRAS, kBET, GC, and sBEE scores. These are followed by a subplot showing their unweighted averages across cell types together with the dataset-level CiLISI and PCR scores. CiLISI and PCR, which are set apart by whitespace, report a single score for the entire scenario rather than cell-type-specific scores. Scores range from 0 to 1; higher values indicate better integration.

We next considered an incomplete-correction scenario involving a single cell type present in three batches. After integration, two batches are well mixed, while the third remains clearly separated (Scenario B, Fig. 3). sBEE (0.17), iLISI (0.25), and CiLISI (0.25) correctly indicate incomplete integration. In contrast, ASW and GC produce overly optimistic scores of 0.80 and 1.00, respectively. kBET yields a score of 0.00, failing to reflect the partial correction, while BRAS (0.43) and PCR (0.42) also overestimate the integration quality.

**Fig. 3:**
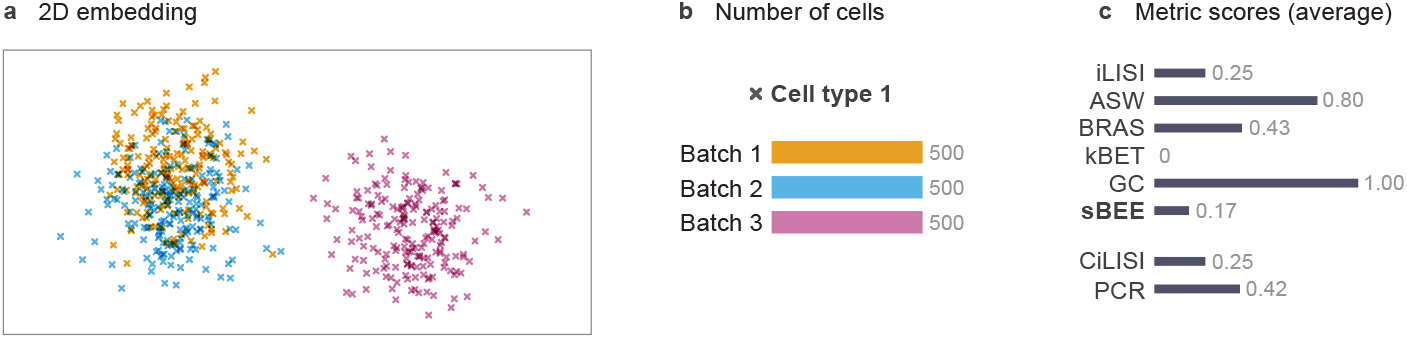
a) Simulated 2D embedding for Scenario B (incomplete batch correction). b) Batch composition, where colors represent batches. c) Metric scores for the single cell type. The dataset-level CiLISI and PCR scores are set apart from the other metrics by whitespace. Scores range from 0 to 1; higher values indicate better integration.

We next assessed scenarios in which batch composition and cell-type coverage complicate the evaluation of batch mixing. We began with a cell-type-specific batch imbalance scenario in which batch effects were equally strong across cell types, but batch 2 contained substantially fewer cells from cell types 2 and 3. Accordingly, scores should be uniformly low across all cell types. sBEE, iLISI, kBET, and CiLISI meet this expectation (Scenario C, Fig. 4), whereas GC (1.00, 0.83, 1.00), ASW (0.48–0.49), and PCR (0.78) substantially overestimate integration quality. BRAS (0.23, 0.33, 0.21) also overestimates integration quality and fails to produce consistent scores across cell types despite the uniform strength of the batch effect. Notably, cell type 2, one of the two imbalanced cell types, receives the highest score. This inconsistency indicates that BRAS is sensitive to cell-type composition.

**Fig. 4:**
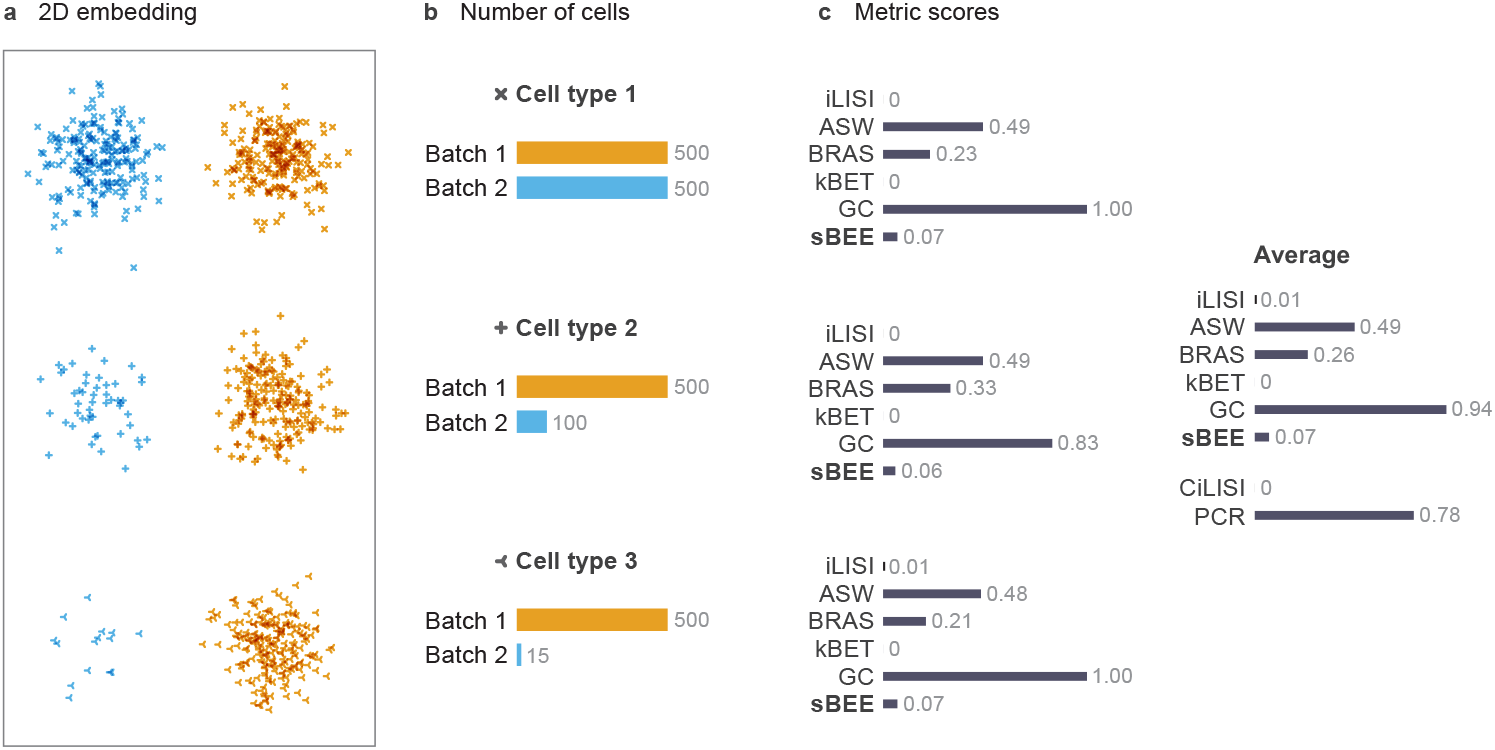
a) Simulated 2D embedding for Scenario C (cell-type-specific batch imbalance). b) Batch composition of each cell type, where colors represent batches. c) Metric scores. Each cell type has a separate subplot showing its iLISI, ASW, BRAS, kBET, GC, and sBEE scores. These are followed by a subplot showing their macro-average across cell types together with the dataset-level CiLISI and PCR scores. CiLISI and PCR, which are set apart by whitespace, report a single score for the entire scenario rather than cell-type-specific scores. Scores range from 0 to 1; higher values indicate better integration.

A complementary scenario in which no cell types are shared between batches is shown in Supplementary Fig. S1 (Scenario D). sBEE, kBET, and GC correctly assign perfect scores, as cross-batch mixing is neither possible nor expected in this setting. ASW and BRAS are undefined because they require multiple batches per cell type. iLISI assigns a score of 0 to both cell types, CiLISI yields 0, and PCR reports 0.29.

These metrics therefore incorrectly penalize the integration for the absence of cross-batch mixing, even though such mixing is impossible when the batches contain different cell types.

We then considered a batch-proportion scenario in which batch separation varies by cell type, with strong separation in cell type 1 and equally mild separation in cell types 2 and 3 despite their different batch proportions (Scenario E). sBEE captures this pattern (0.06 vs. 0.66–0.71; Fig. 5) and remains robust to the imbalance in cell type 3. In contrast, iLISI and kBET are sensitive to this imbalance, while ASW, GC, and PCR overestimate mixing and fail to resolve the cell-type-specific differences. BRAS overestimates mixing under strong separation (cell type 1), and CiLISI underestimates the overall integration quality (0.27).

**Fig. 5:**
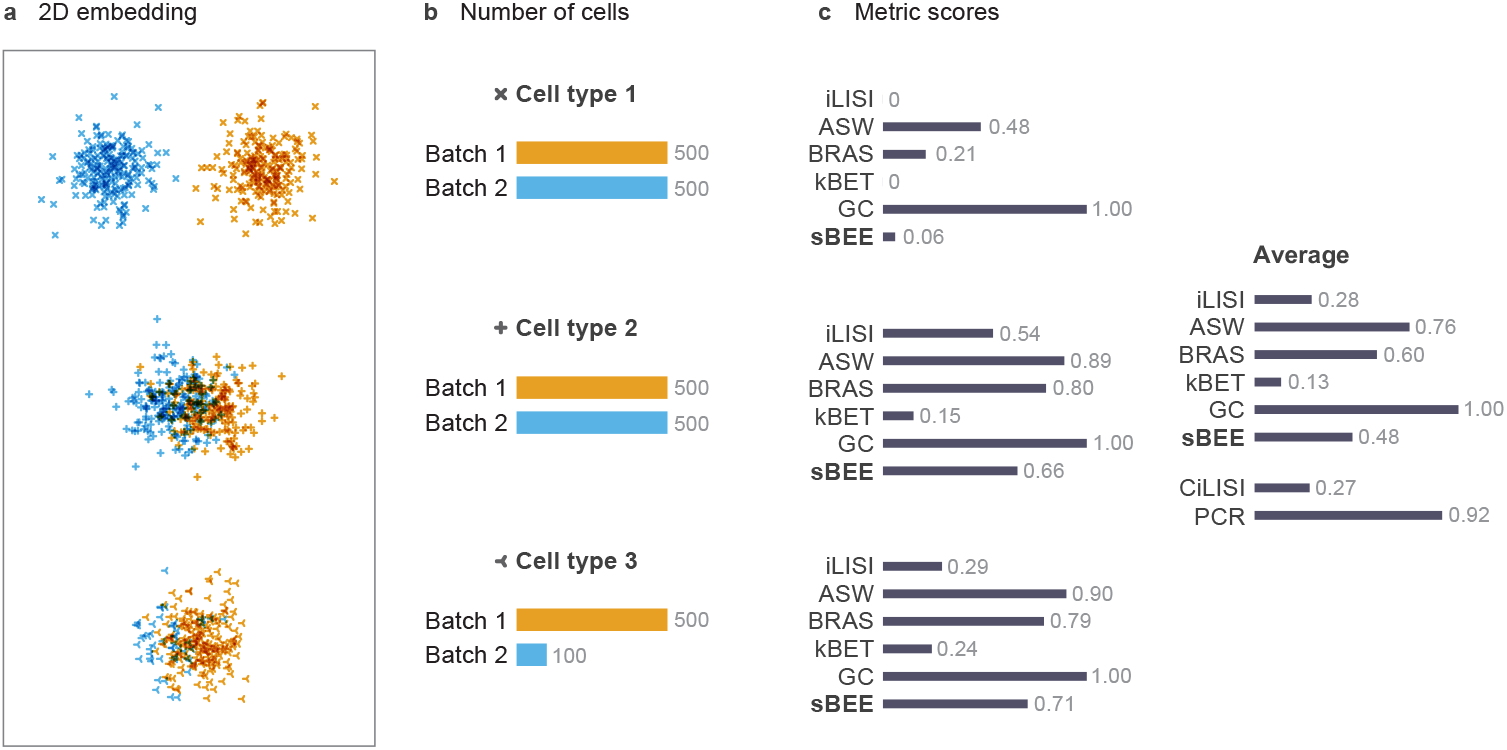
a) Simulated 2D embedding for Scenario E (robustness to batch-proportion imbalance). b) Batch composition of each cell type, where colors represent batches. c) Metric scores. Each cell type has a separate subplot showing its iLISI, ASW, BRAS, kBET, GC, and sBEE scores. These are followed by a subplot showing their macro-average across cell types together with the dataset-level CiLISI and PCR scores. CiLISI and PCR, which are set apart by whitespace, report a single score for the entire scenario rather than cell-type-specific scores. Scores range from 0 to 1; higher values indicate better integration.

To assess whether the metrics can resolve graded differences in batch-effect severity across cell types, we constructed Scenario F (Fig. 6). sBEE accurately captures this gradient. iLISI preserves the ordering but compresses moderate and strong separation into near-zero scores, while kBET similarly fails to distinguish between these levels. ASW, GC, and PCR substantially overestimate integration quality and fail to capture the gradient in batch-effect severity. BRAS overestimates integration quality under moderate separation (cell type 2), while CiLISI reports an overall score of 0.29, indicating intermediate integration quality.

**Fig. 6:**
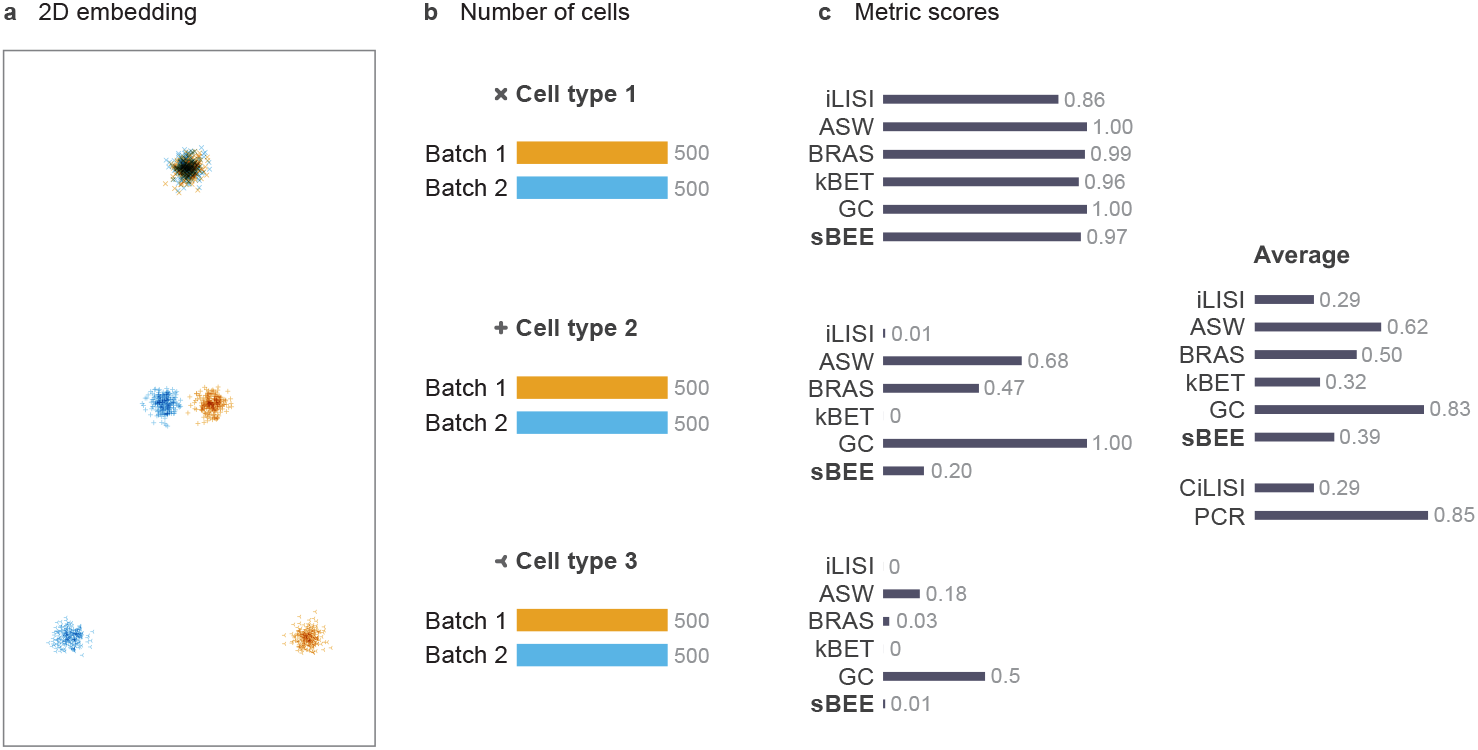
a) Simulated 2D embedding for Scenario F (graded batch effects). b) Batch composition of each cell type, where colors represent batches. c) Metric scores. Each cell type has a separate subplot showing its iLISI, ASW, BRAS, kBET, GC, and sBEE scores. These are followed by a subplot showing their macro-average across cell types together with the dataset-level CiLISI and PCR scores. CiLISI and PCR, which are set apart by whitespace, report a single score for the entire scenario rather than cell-type-specific scores. Scores range from 0 to 1; higher values indicate better integration.

We then considered Scenario G (Supplementary Fig. S3), in which the overlap between batches for cell type 2 increases progressively, while cell type 1 remains completely separated by batch. For cell type 1, ASW and GC nevertheless assign optimistic scores despite the complete absence of batch mixing, indicating limited sensitivity to the persistent separation. Turning to cell type 2, sBEE increases monotonically with increasing overlap and clearly distinguishes the successive levels of batch mixing. iLISI and kBET also increase with overlap, but their scores remain clustered near the lower end at low overlap and therefore distinguish these settings less clearly. In contrast, GC remains saturated at 1.00 across all settings and therefore fails to capture the progressive change. ASW and PCR also increase with overlap but assign relatively high scores at low and intermediate mixing levels. BRAS similarly returns a non-negligible score of 0.23 when there is no overlap between batches for cell type 2, although it subsequently increases monotonically. For cell type 2, CiLISI responds weakly at low overlap, increasing only from 0.02 to 0.08 by 50% overlap, followed by sharper increases to 0.26 and 0.42 at 75% and 100% overlap, respectively. Its value of only 0.42 at complete overlap suggests that the metric does not fully reflect the perfect batch mixing achieved for this cell type.

To assess whether the metrics capture differences in the severity of complete batch separation, we compared two configurations in Scenario H (Supplementary Fig. S4a,b). sBEE clearly captures this distinction, decreasing from 0.27 in the closer configuration to 0.01 in the more distant configuration. In contrast, iLISI, CiLISI, and kBET assign near-zero scores to both configurations and therefore fail to distinguish their relative severity. ASW and PCR decrease as the separation increases but assign overly favorable scores to the closer configuration despite the complete absence of batch mixing. GC is similarly optimistic, returning scores of 1.00 and 0.50 for the closer and more distant configurations, respectively. BRAS also substantially overestimates integration quality in the closer configuration.

We next examined whether the metrics respond appropriately to a more subtle increase in cross-batch overlap in Scenario I (Supplementary Fig. S4c,d). sBEE distinguishes the two levels, increasing from 0.11 to 0.21 as the overlap increases. iLISI and CiLISI show the same directional change but exhibit a smaller difference between the two configurations. kBET changes only marginally, while GC remains saturated at 1.00 and therefore fails to reflect the increase in overlap. ASW, BRAS, and PCR also increase with overlap but assign relatively high scores, particularly in the lower-overlap configuration, thereby overstating the degree of batch mixing.

To further evaluate metric sensitivity to progressively increasing batch overlap, Scenario J (Supplementary Fig. S6) uses a concentric configuration with increasing spread. sBEE increases monotonically with spread, clearly reflecting the improvement in batch mixing. iLISI, kBET, and CiLISI show the same general trend. ASW also increases but assigns an overly optimistic score at the smallest spread, despite the limited overlap between batches. BRAS remains nearly saturated and even decreases slightly across the progression (0.99, 0.99, and 0.97), failing to reflect the increasing overlap. GC and PCR are entirely insensitive to this change, remaining fixed at 1.00 throughout.

Scenarios K and L (Supplementary Fig. S6) extend this evaluation to off-center configurations in which geometric constraints prevent complete batch mixing, even at large spread values. In both scenarios, sBEE increases with spread but appropriately remains below its maximum value. It also assigns consistently lower scores to Scenario L than to Scenario K, capturing the stronger residual separation produced by the two off-center batches. iLISI and CiLISI exhibit similar qualitative behavior. In contrast, kBET remains overly conservative and underestimates batch mixing across all configurations, while ASW, BRAS, GC, and PCR systematically assign scores that are too high for the degree of overlap achieved.

#### Robustness to parameter choices

We assessed the sensitivity of sBEE to its two parameters, the separation-component scale *σ* and the neighborhood size *k* (Supplementary Fig. S8). Across the five scenarios examined (C, E, F, H, and J), sBEE scores were essentially invariant to *k* over the range 30–120, consistent with the per-cell-type cap imposed on the neighborhood size. In contrast, *σ* affects the absolute scale of the scores, although the relative ordering of configurations and their qualitative interpretations remain stable across a broad range of values. At the default setting of *σ* = 0.15, configurations expected to receive low scores, such as the cell-type-specific batch-size imbalance in Scenario C and the strongly separated cell type in Scenario F, remain appropriately low, while well-mixed configurations retain high scores. We therefore selected *σ* = 0.15 because it preserves low scores for poorly integrated configurations while maintaining clear discrimination among settings with different degrees of residual batch separation. Larger values of *σ* systematically increase the scores and compress the differences among configurations.

### 2.2 Evaluation across real data

We evaluated the metrics under two evaluation settings. First, using the human pancreas and lung atlases [15], we applied seven integration methods: Scanorama [12], MNN [6], ComBat [4], scANVI [28], scVI [9], Harmony [7], and scGen [14]. We then assessed whether each metric consistently ranked the integrated datasets above the corresponding unintegrated baseline. Second, for the BMMC (NeurIPS 2021) [29] and HBCA [30] datasets, we evaluated liam-integrated [31] representations corresponding to progressively increasing levels of batch removal and examined whether metric scores increased monotonically across these ordered integration stages. In both settings, a reliable metric should assign higher scores as integration quality improves.

#### 2.2.1 Metric Scores for Integrated versus Unintegrated Data

##### Pancreas benchmark

In the human pancreas atlas [15], we assessed whether each metric assigned higher scores to the representations produced by the integration methods than to the unintegrated data. Red bars indicate cases in which the metric score for an integrated representation is lower than or equal to the corresponding score for the unintegrated data (Fig. 7). iLISI, CiLISI, BRAS, kBET, PCR, and sBEE assigned higher scores to every integrated representation than to the unintegrated baseline.

**Fig. 7:**
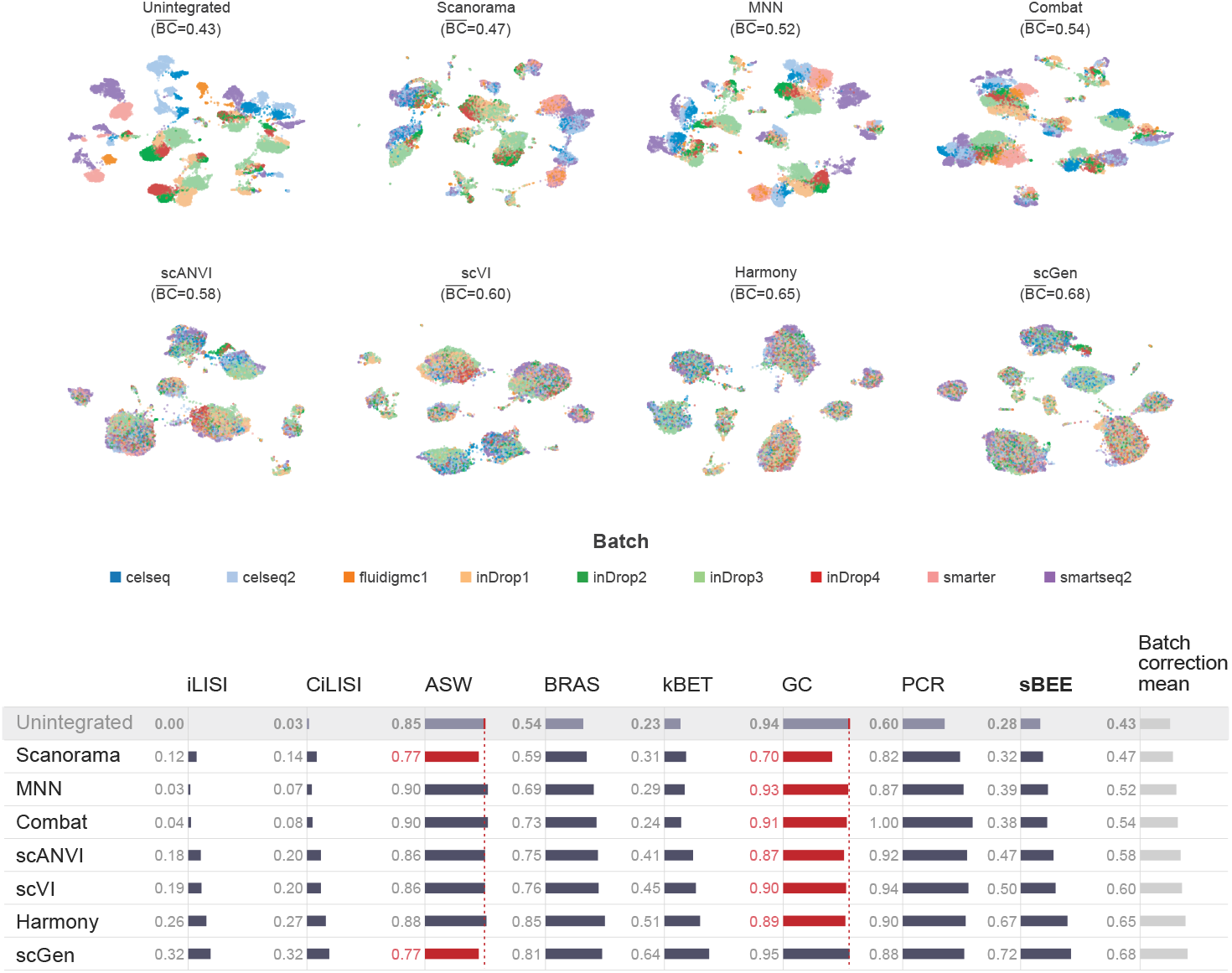
Pancreas [15] benchmark. The upper panel shows UMAPs of the unintegrated data and the outputs of the seven integration methods, ordered by mean batch correction score. The lower panel shows rankings of the integration methods; rows are ordered by mean batch correction score, except for the unintegrated row, which is fixed at the top. The batch correction mean is calculated as the average of the individual batch correction metrics. Red bars indicate method–metric pairs whose score is lower than or equal to the corresponding unintegrated score for that metric.

**Fig. 8:**
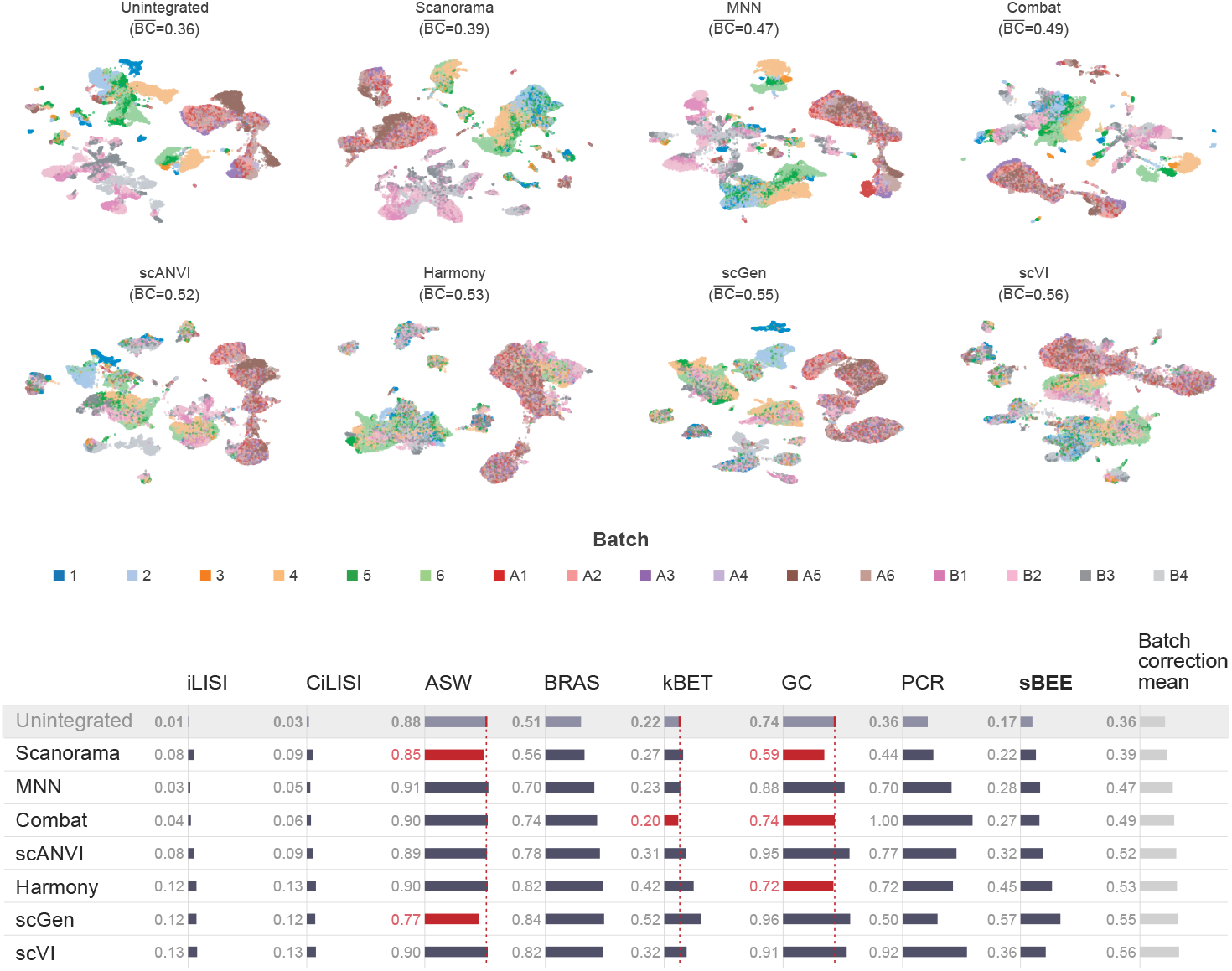
Lung [15] benchmark. The upper panel shows UMAPs of the unintegrated data and the outputs of the seven integration methods, ordered by mean batch correction score. The lower panel shows rankings of the integration methods; rows are ordered by mean batch correction score, except for the unintegrated row, which is fixed at the top. The batch correction mean is calculated as the average of the individual batch correction metrics. Red bars indicate method–metric pairs whose score is less than or equal to the corresponding unintegrated score for that metric.

However, the magnitude of the improvement varied among these metrics. For example, kBET only marginally distinguished ComBat [4] from the unintegrated baseline (0.24 versus 0.23), while iLISI and CiLISI assigned relatively low scores to MNN [6] and ComBat [4]. sBEE consistently distinguished all integration methods from the baseline, increasing from 0.28 for the unintegrated data to values ranging from 0.32 to 0.72 after integration. It assigned its highest scores to Harmony [7] (0.67) and scGen [14] (0.72), which also received the two highest mean batch-correction scores. ASW and GC, however, frequently failed to rank the integrated representations above the unintegrated baseline. ASW assigned lower scores to Scanorama [12] and scGen [14] than to the unintegrated data (0.77 and 0.77 versus 0.85), despite the increased batch mixing visible in their embeddings. GC showed an even stronger baseline bias, assigning a score of 0.94 to the unintegrated data. Consequently, it ranked six of the seven integrated representations below the unintegrated one. These results indicate that the high baseline scores produced by ASW and particularly GC limit their ability to detect improvements in batch mixing in the pancreas dataset.

##### Lung benchmark

We conducted a similar assessment in the human lung atlas [15]. iLISI, CiLISI, BRAS, PCR, and sBEE assigned higher scores to every integrated representation than to the unintegrated data. sBEE increased from 0.17 for the unintegrated data to values ranging from 0.22 to 0.57 after integration. It assigned its highest score to scGen [14] (0.57), followed by Harmony [7] (0.45) and scVI [9] (0.36). kBET ranked six of the seven integrated representations above the baseline but assigned ComBat [4] a lower score than the unintegrated data (0.20 versus 0.22). ASW failed to identify improvements for Scanorama [12] and scGen [14], assigning them scores of 0.85 and 0.77, respectively, compared with 0.88 for the unintegrated data. GC also failed for Scanorama [12], ComBat [4], and Harmony [7], with scores of 0.59, 0.74, and 0.72, respectively, compared with the unintegrated score of 0.74. Thus, as in the pancreas benchmark, the high baseline values produced by ASW and GC limited their ability to recognize improved batch mixing, whereas sBEE consistently ranked all integrated representations above the unintegrated baseline.

#### 2.2.2 Cell-Type-Level Evaluation of Partial Batch Correction

To gain more detailed insight into metric behavior on real data, we examined alpha cells from the unintegrated and MNN-integrated pancreas datasets [6]. Alpha cells were selected because they were the most abundant cell type in the pancreas dataset, accounting for more than 33.5% of all cells, and because the metrics produced strongly discordant scores for this cell type. Within the alpha-cell subset (Fig. 9), sBEE increased from 0.14 for the unintegrated data to 0.33 after MNN integration. iLISI and CiLISI also increased from 0.01 to 0.05, but both remained close to zero. Although iLISI and CiLISI consistently ranked the integrated representations above the unintegrated baseline in the preceding dataset-level evaluations, the compressed range observed here indicates limited sensitivity to the mixing level. ASW increased from 0.82 to 0.93, and BRAS increased from 0.46 to 0.70. In contrast, kBET remained at 0 and GC remained at 0.96 for both representations.

**Fig. 9:**
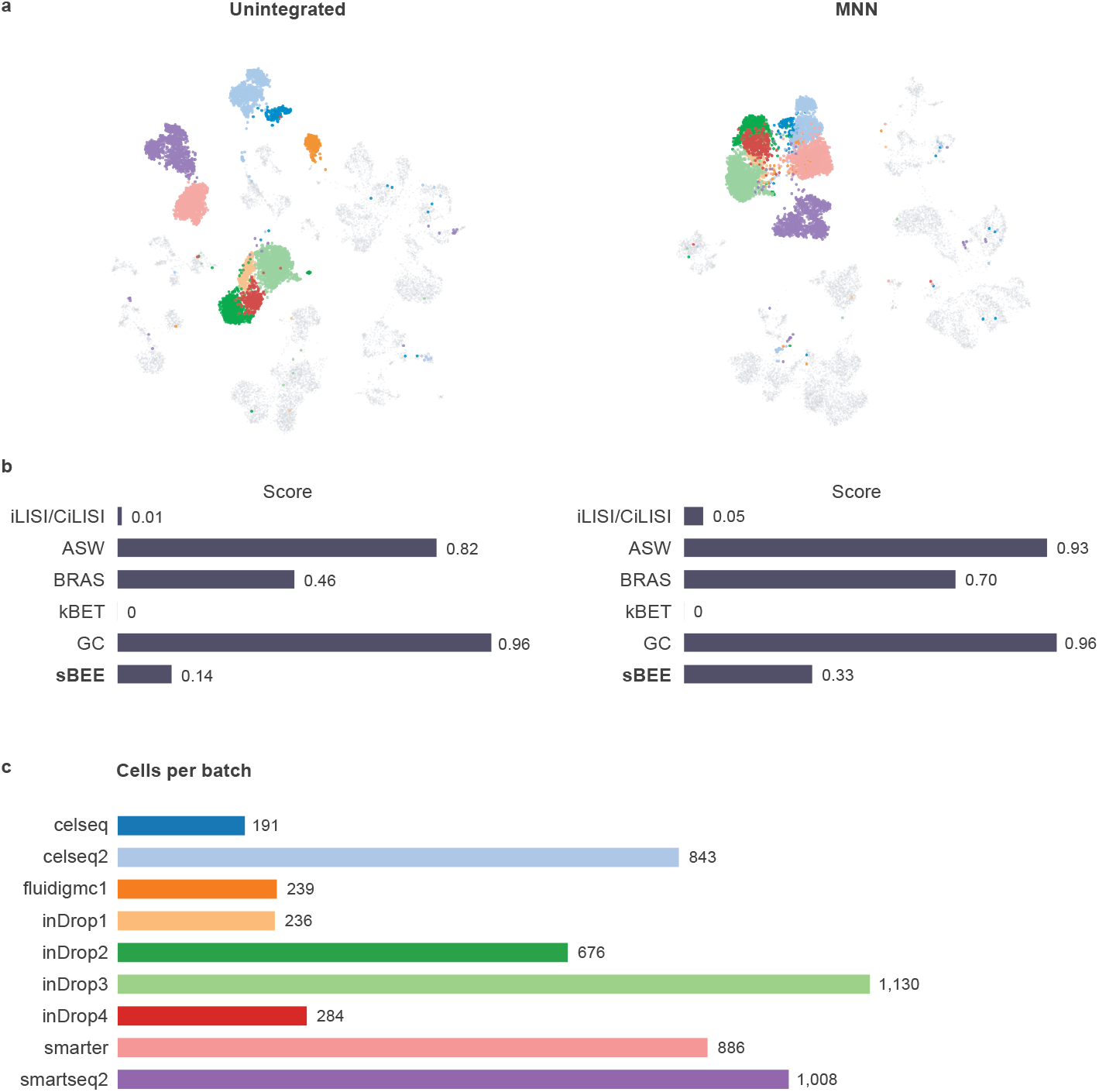
Batch mixing of alpha cells before and after MNN integration. a) UMAP embeddings of the unintegrated and MNN-integrated pancreas datasets. Alpha cells are colored by batch, while all other cell types are shown in grey. b) Batch-correction metrics computed for alpha cells. c) Number of alpha cells that exist in each batch.

The corresponding UMAP embeddings (Fig. 9a) indicate that the unintegrated and MNN-integrated alpha-cell representations should receive different batch-correction scores. In the unintegrated data, the batches form clearly separated groups. Following MNN integration, these batch-specific groups move closer together but remain incompletely mixed. Thus, the near-zero iLISI and CiLISI scores are consistent with the limited sensitivity under strong residual separation observed in the simulated scenarios. ASW and BRAS, in contrast, appear overly optimistic, assigning relatively high scores even when cells remain visibly clustered by batch. kBET does not distinguish the two representations under its heuristic neighborhood-size selection, while GC also remains unchanged. Among the evaluated metrics, sBEE most closely reflects the qualitative structure of the embeddings by assigning a low score to the clearly separated unintegrated data and a higher, but still moderate, score to the partially corrected MNN representation. The same qualitative behavior was also observed for beta cells (Supplementary Fig. S7).

#### 2.2.3 Sensitivity to Ordered Levels of Batch Correction

Unlike the pancreas and lung benchmarks, in which the outputs of different integration methods do not have an explicit ground-truth ordering, the BMMC and HBCA benchmarks provide representations with predefined, progressively increasing levels of batch removal. This design allows a direct evaluation of whether a metric responds consistently to gradual improvements in integration quality. For BMMC, we used four ordered integration stages provided by [24], generated using liam’s [31] batch-adversarial training strategy, in which a tunable scaling parameter *α* controls the strength with which batch-associated variation is penalized and removed from the learned embedding: unintegrated (no batch correction), suboptimal (low *α*, only partially removing batch effects), effective (moderate *α*, substantially reducing batch effects while preserving biological variation), and optimized (high *α*, maximizing batch mixing) (Fig. 10). Because *α* directly and monotonically controls the magnitude of the adversarial penalty during training, increasing *α* provides a principled axis for generating embeddings with progressively stronger batch removal, independent of any particular evaluation metric. The UMAP embeddings were visualized by batch to show how biological structure was retained while batch-associated separation was progressively reduced across these four stages. BMMC presents an additional challenge

**Fig. 10:**
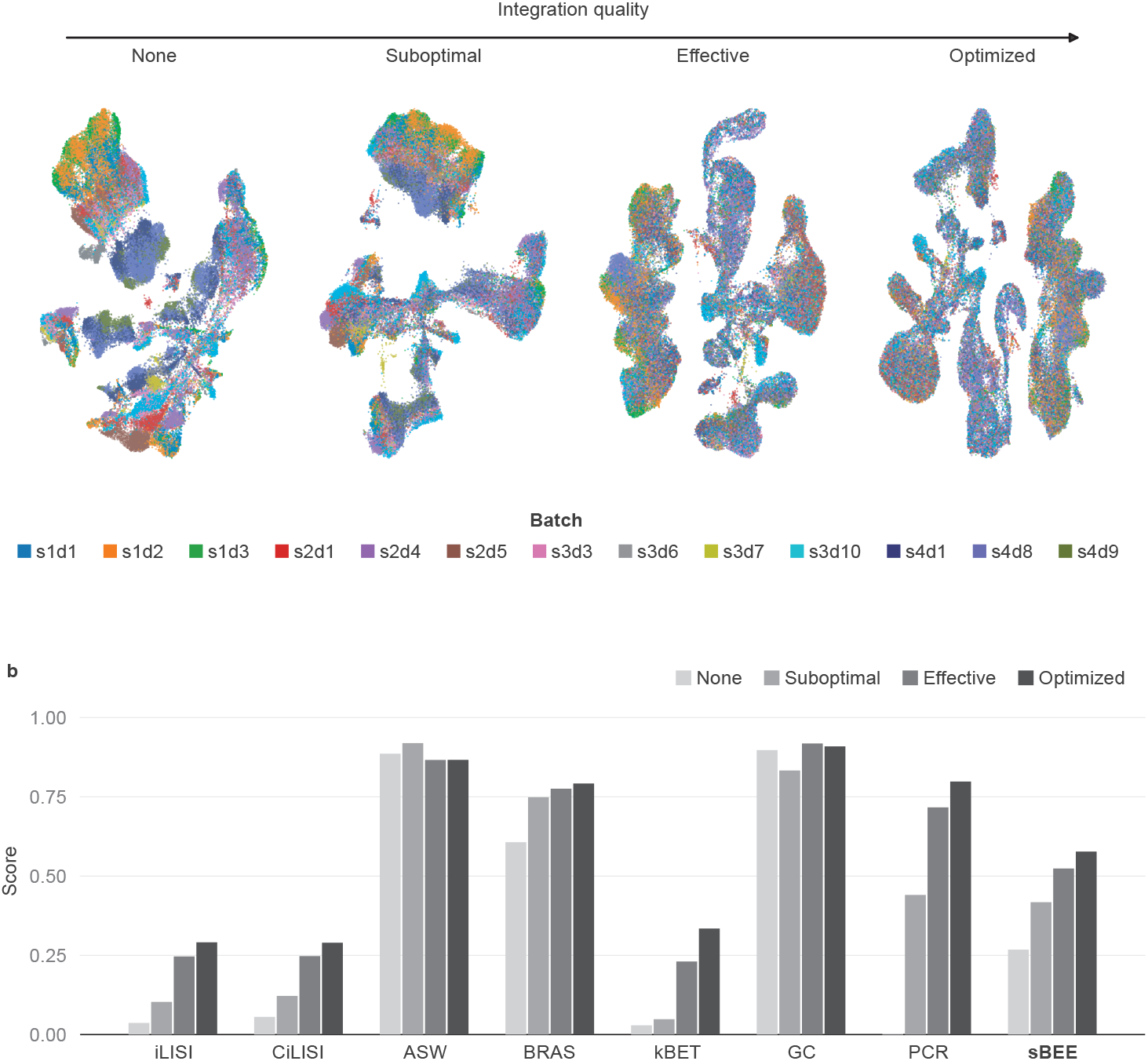
BMMC benchmark (NeurIPS 2021) [29]. a) UMAP embeddings of the four integration stages: unintegrated, suboptimal, effective, and optimized. b) Batch-correction metric scores across the four integration stages.

because its batch structure is nested, with variation arising from both donors and collection sites. A metric was expected to increase at each successive stage despite this multilevel batch structure. For HBCA, we similarly used three ordered integration stages from [24], representing progressively stronger batch correction obtained via the same *α*-tuning strategy (Fig. 11).

**Fig. 11:**
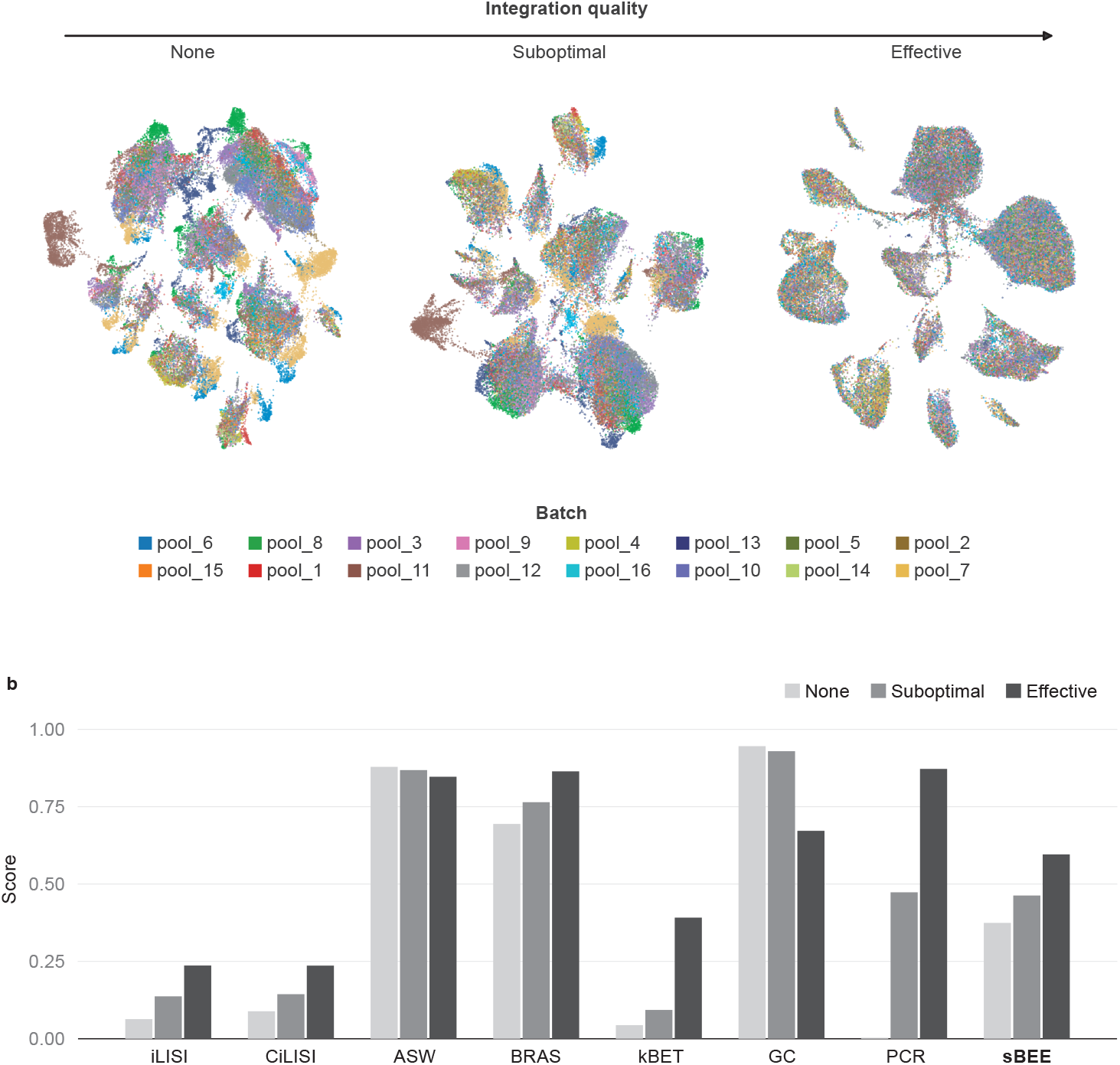
HBCA [30] dataset. a) UMAP embeddings of the four integration stages: unintegrated, suboptimal, effective, and optimized. b) Batch-correction metric scores across the four integration stages/

Across the ordered integration stages in BMMC and HBCA, sBEE, iLISI, CiLISI, BRAS, kBET, and PCR increased monotonically with progressively stronger batch removal (Figs. 10 and 11). In contrast, ASW and GC failed to follow the expected stage-wise improvement in integration quality. Among the metrics that preserved the correct ordering, BRAS assigned relatively high scores to the unintegrated representations, with values of 0.61 for BMMC and 0.69 for HBCA. Thus, although BRAS captured the relative improvement across stages, it overestimated the absolute integration quality of the unintegrated data.

### 2.3 Recurring failure modes of existing batch-mixing metrics

The controlled simulations reveal several systematic failure modes in the existing batch-mixing metrics (Table 1). The real-data analyses corroborated several of these patterns, indicating that the observed limitations are not restricted to synthetic settings.

**Table 1:** Failure modes observed for the evaluated batch-mixing metrics. ✗ indicates that the corresponding failure mode was observed, while a blank cell indicates that it was not observed in the evaluated scenarios. None of the listed failure modes was observed for sBEE.

| Metric | Partial batch coverage | Batch imbalance sensitivity | Separation sensitivity | Over-optimism | Directional failure | Structural insensitivity | Masking cell-type specific effects |
| --- | --- | --- | --- | --- | --- | --- | --- |
| **sBEE** |  |  |  |  |  |  |  |
| iLISI | **X**<br>A,D | **X**<br>E | **X**<br>F,H |  |  |  |  |
| CiLISI | **X**<br>A,D |  | **X**<br>H |  |  |  |  |
| kBET |  | **X**<br>E | **X**<br>B,F,H,I |  |  |  |  |
| ASW |  |  |  | **X**<br>B,C,E,F<br>G,H,I,K,L |  |  |  |
| BRAS |  |  |  | **X**<br>B,C,E,F<br>G,H,I,K,L | **X**<br>J |  |  |
| GC |  |  |  |  |  | **X**<br>B,C,E,F<br>G,H,I,J<br>K,L |  |
| PCR | **X**<br>A,D |  |  | **X**<br>B,C,E,F<br>G,H,I,J<br>K,L |  |  | **X**<br>E,F,G |

#### Partial batch coverage

iLISI, CiLISI and PCR incorrectly penalize cell types that appear in only a subset of batches (scenarios A and D). All three metrics conflate data-intrinsic batch absence with integration failure.

#### Batch imbalance sensitivity

iLISI and kBET assign different scores to cell types with comparable residual batch separation but unequal batch sizes, making their outputs sensitive to batch composition rather than integration quality alone (scenario E).

#### Separation sensitivity

iLISI and kBET fail to distinguish moderate from strong batch separation, collapsing to near-zero values (scenarios F, H); CiLISI shows the same collapse when batches are clearly separated (scenario H). kBET additionally exhibits a binary response in partial correction settings, collapsing to zero even when the majority of cells are well mixed (scenario B), and assigns nearly identical scores to low and high overlap (scenario I).

#### Overestimation of integration quality

ASW and BRAS frequently assign high scores despite substantial residual batch effects, across partial correction, cell-typespecific batch imbalance, graded effects, and non-standard batch geometries (scenarios B, C, E, F, G, H, I, K, L). PCR overestimates integration quality across nearly all of these settings as well (scenarios B, C, E, F, G, H, I, J, K, L).

#### Directional failure

In concentric batch configurations (scenario J), BRAS fails to increase as batch mixing improves; rather than rising with overlap, its scores remain essentially flat and edge slightly downward (0.99, 0.99, 0.97). Although the magnitude of this reversal is small, the absence of the expected upward trend indicates that BRAS does not reliably track mixing when one batch forms a ring-like structure surrounding another in the embedding space.

#### Structural insensitivity

GC measures whether cells of the same type form a single connected component in the *k*-nearest neighbor graph, a condition that is satisfied by almost any embedding after integration. As a result, GC outputs scores close to or equal to 1.00 in nearly every scenario regardless of actual batch mixing quality (scenarios B, C, E, F, G, H, I, J, K, L), scoring correctly only when a high score is genuinely warranted (scenarios A and D) or when the graph is genuinely disconnected.

#### Masking of cell-type-specific effects

PCR aggregates batch variance across the entire dataset rather than evaluating cell types individually, obscuring cell-type-specific differences in integration quality (scenarios E, F, and G).

The real-data results supported several of these simulation-derived failure modes. In the pancreas alpha-cell analysis, iLISI and CiLISI remained close to zero for both unintegrated and MNN-integrated data, consistent with the separation sensitivity observed in the simulations. ASW and BRAS assigned relatively high scores despite visible batch-associated clustering, consistent with their over-optimism in the controlled scenarios. kBET did not distinguish the two embeddings with the neighborhood size selected by its heuristic, and GC remained unchanged at a high score, consistent with its structural insensitivity. Similar qualitative behavior was also observed for beta cells (Supplementary Fig. S7). These observations suggest that the recurring failure modes identified in controlled simulations can also affect metric behavior on real datasets.

In contrast, sBEE consistently reflected the true degree of batch mixing in the controlled simulations and showed concordant behavior in the real-data analyses, providing a more reliable evaluation of integration quality.

## 3 Discussion

Our analysis shows that widely used batch-mixing metrics, despite their broad adoption, can misrepresent integration quality under common but non-trivial conditions. Because benchmarking studies typically report a single aggregate score for each method, these failure modes can be easily masked. A metric may produce plausible overall rankings while remaining sensitive to data characteristics such as data-intrinsic batch absence, batch-size imbalance, or non-globular cluster geometry. By isolating each challenge in a controlled scenario with a known expected outcome, we were able to identify these failure modes independently of dataset-specific variation.

The observed failure modes largely reflect differences in metric design. Neighborhood-composition metrics such as iLISI and CiLISI penalize structurally missing cell type and batch combinations because a batch in which a cell type is absent cannot appear in the local neighborhoods of that cell type, regardless of integration quality. These metrics also tend to collapse toward zero when batches are clearly separated, limiting their ability to distinguish different degrees of residual separation. kBET shows a related loss of dynamic range because its hypothesis-testing framework can reject local batch mixing even when most cells are reasonably well integrated. Silhouette-based metrics such as ASW and BRAS are strongly influenced by cluster geometry and can produce overly optimistic scores for non-globular or concentric clusters. GC is structurally insensitive to within-component batch separation. Cells of the same type may remain connected in the graph despite poor batch mixing, causing GC to report near-maximal scores except when the graph is genuinely fragmented. PCR summarizes batch-associated variance globally using a linear decomposition, which can lead it to overestimate integration quality while obscuring cell-type-specific residual effects.

sBEE was designed to avoid these failure modes by combining two complementary views of batch mixing within each cell type: local neighborhood composition, measured by the agreement between neighborhood and global batch proportions, and cross-batch distances, measured by the ratio of intra-to inter-batch distances. Jointly evaluating these signals at the cell-type level allows sBEE to provide reliable scores in settings where single-signal metrics fail. It does not penalize cell types that are absent from some batches, retains dynamic range across different degrees of separation, and remains robust to batch-size imbalance because batches contribute equally within each cell type and the cell-type-specific scores are macro-averaged. Across the pancreas and lung datasets, sBEE behaved as expected across integration methods. For the BMMC and HBCA datasets, it also reflected progressively stronger levels of integration.

One limitation in our study is that the simulated scenarios are generated as Gaussian clusters in a two-dimensional latent space and lifted to higher dimensions through an orthonormal projection. Although this design enables precise control over cluster geometry and batch overlap, it does not reproduce the full complexity of real scRNA-seq count data. More realistic count-based scRNA-seq simulators could better reproduce the distributional properties of real expression data, but, as mentioned in the Methods, they provide limited control over cluster geometry and batch coverage. Our simulation design therefore prioritizes experimental control over generative realism, allowing individual challenges to be isolated and evaluated against predefined outcomes. Our real-data evaluation is further limited to four scRNA-seq datasets and the integration methods considered in this study. Evaluation across additional tissues, experimental technologies, batch structures, and integration methods will be necessary to establish the generalizability of the findings.

sBEE includes a sensitivity parameter, *σ*, and, like other neighborhood-based metrics, depends on the neighborhood size, *k*. A sensitivity analysis across scenarios (Supplementary Fig. S8) shows that the qualitative conclusions are preserved over a broad range of *σ* values and that the scores are essentially invariant to *k*. We selected *σ* = 0.15 because it keeps scores low when integration is poor while preserving discrimination among scenarios with different degrees of separation. For each cell type *c*, the effective neighborhood size is defined as *k_c_* = min(*k,* |*C_c_*| − 1), where |*C_c_*| is the number of cells of type *c*. The neighborhood size therefore adapts to the number of available cells in each cell type and never extends beyond cell-type boundaries. Consequently, neighborhood-composition estimates for very small cell types are based on few cells and may therefore be noisier.

Finally, sBEE evaluates batch mixing only and is intended to be used alongside, rather than in place of, biological-conservation metrics. A metric that rewards mixing without considering biological structure can be trivially maximized through overcorrection. Several extensions follow naturally from this work. Although the definition of *k_c_* makes the neighborhood size adaptive across cell types, fully data-driven neighborhood schemes could eliminate dependence on a fixed default value of *k* altogether. Another promising direction is to combine sBEE with a biological-conservation term to obtain a composite score and evaluate it on established integration benchmarks.

## 4 Conclusion

Through a systematic evaluation of batch-mixing metrics across simulated integration scenarios, we identified several recurring failure modes in existing approaches. Motivated by these observations, we introduced sBEE, a metric that evaluates batch mixing within each cell type from two complementary views: local neighborhood composition and cross-batch distances. Across twelve scenarios, sBEE consistently produced scores that reflected the true degree of batch mixing, correctly handling batch-specific cell types, partial batch coverage, batch-size imbalance, non-globular geometries, partial correction, and progressive mixing. In contrast, every existing metric failed in at least one scenario. sBEE also handled imbalanced batch compositions, remained insensitive to the neighborhood size *k*, and produced consistent verdicts across a broad range of its scale parameter *σ*. On the real pancreas and lung datasets it behaved correctly across integration methods and, for BMMC and HBCA, across progressively stronger integration stages.

Together, our results provide both a systematic characterization of the strengths and limitations of current batch-mixing metrics and a practical alternative that is robust across a broad range of integration settings. We therefore recommend sBEE as a primary metric for evaluating single-cell batch mixing, particularly when partial batch coverage or non-standard embedding geometries are present, and suggest using it alongside biological-conservation metrics rather than in isolation. Future work includes combining sBEE with a biological-conservation metric into a single composite score for routine use.

## 5 Methods

We first describe the simulated scenarios designed to isolate key integration challenges and the real datasets used for evaluation. We then describe the preprocessing pipeline, detail the existing metrics used for benchmarking, and finally introduce the proposed metric.

### 5.1 Simulated Evaluation Scenarios

We designed a set of simulated scenarios to evaluate batch integration metrics under controlled conditions. Each scenario isolates a specific integration challenge, including batch imbalance, partial batch coverage (i.e., cell types represented in only a subset of batches), non-globular cluster geometry, and varying batch effect strengths. We did not use count-based scRNA-seq simulators such as Splatter [32] because they provide limited control over cluster geometry and batch coverage. Instead, we generated data directly in a two-dimensional latent space and applied an orthonormal projection to obtain 2000-dimensional observations. In all scenarios, cell types correspond to Gaussian clusters in the two-dimensional latent space, while batch effects are introduced by controlled shifts of cluster centers between batches. The orthonormal projection preserves these geometric relationships in the high-dimensional space. Detailed parameters for each scenario are provided in Supplementary Table 1. Both simulated and real datasets are preprocessed as described below before metric evaluation. For each scenario, we define the expected behavior of an ideal integration metric. Any deviation from this expected behavior is considered a failure. A summary of all scenarios is provided in Table 2.

**Table 2:**
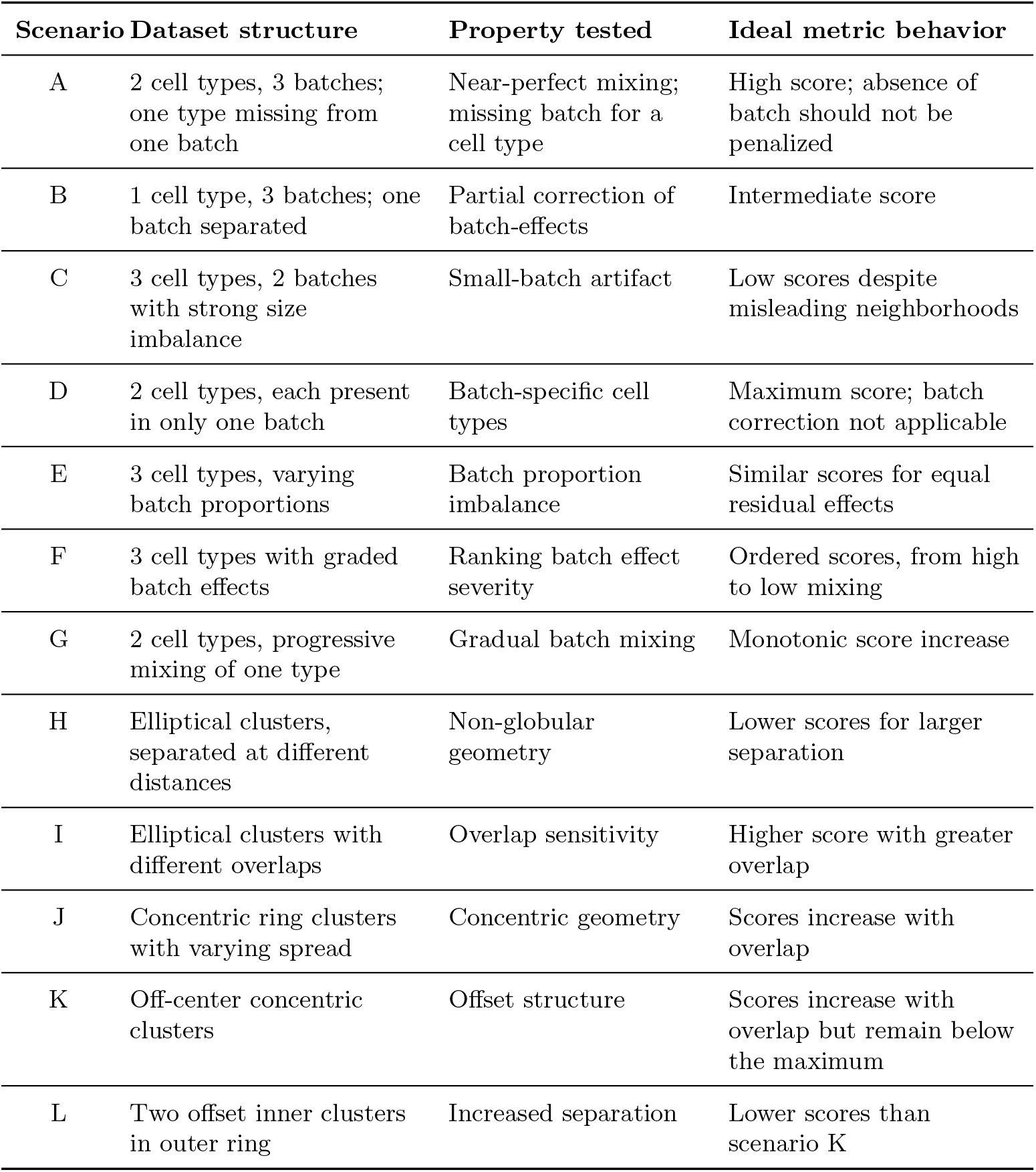
Summary of simulated evaluation scenarios used to evaluate batch integration metrics.

| Scenario | Dataset structure | Property tested | Ideal metric behavior |
| --- | --- | --- | --- |
| A | 2 cell types, 3 batches; one type missing from one batch | Near-perfect mixing; missing batch for a cell type | High score; absence of batch should not be penalized |
| B | 1 cell type, 3 batches; one batch separated | Partial correction of batch-effects | Intermediate score |
| C | 3 cell types, 2 batches with strong size imbalance | Small-batch artifact | Low scores despite misleading neighborhoods |
| D | 2 cell types, each present in only one batch | Batch-specific cell types | Maximum score; batch correction not applicable |
| E | 3 cell types, varying batch proportions | Batch proportion imbalance | Similar scores for equal residual effects |
| F | 3 cell types with graded batch effects | Ranking batch effect severity | Ordered scores, from high to low mixing |
| G | 2 cell types, progressive mixing of one type | Gradual batch mixing | Monotonic score increase |
| H | Elliptical clusters, separated at different distances | Non-globular geometry | Lower scores for larger separation |
| I | Elliptical clusters with different overlaps | Overlap sensitivity | Higher score with greater overlap |
| J | Concentric ring clusters with varying spread | Concentric geometry | Scores increase with overlap |
| K | Off-center concentric clusters | Offset structure | Scores increase with overlap but remain below the maximum |
| L | Two offset inner clusters in outer ring | Increased separation | Lower scores than scenario K |

#### Standard Batch Configurations (A–G)

These seven scenarios examine metric behavior under common batch integration challenges, including partial batch coverage (cell types represented in only a subset of batches), incomplete batch correction, batch imbalance, varying batch effect strengths, and progressive batch mixing.

##### A: Near-perfect mixing with a missing batch

The dataset contains two cell types and three batches. For cell type 1, batches 1 and 2 each contain 500 cells, while batch 3 is absent. Cell type 2 is present in all three batches with 500 cells per batch. After integration, batches fully overlap within each cell type, representing near-perfect mixing (Fig. 2). An ideal metric should assign scores close to the maximum for all cell types, recognizing that the absence of batch 3 for cell type 1 is a structural property of the data rather than an integration failure.

##### B: Partial batch correction

The dataset contains one cell type and three batches (500 cells per batch). After integration, two batches are well mixed while the third batch remains separated (Fig. 3). This represents a case where integration has succeeded for the majority of cells but has failed entirely for one batch. An ideal metric should assign an intermediate score.

##### C: Small-batch artifact

The dataset contains three cell types and two batches. Cell type 1 has a balanced 500/500 split, cell type 2 has a 500/100 split, and cell type 3 has a 500/15 split. After integration, all three cell types exhibit the same strong residual batch separation (Fig. 4). Thus, an ideal metric should assign similarly low scores to all three cell types, regardless of the increasing batch-size imbalance. However, as the minority batch becomes smaller, for many integration metrics typical neighborhood sizes inevitably include many cells from the majority batch. Consequently, the local batch composition increasingly resembles the global batch proportions, even though the batches remain well separated.

##### D: Batch-specific cell types

The dataset contains two cell types and two batches, where each cell type is confined to exactly one batch (500 cells per cell type). Since no cell type appears in more than one batch, there is no cross-batch variation to correct, and integration is not applicable (Supplementary Fig. S1). An ideal metric should recognize that the absence of cross-batch neighbors is not a sign of poor integration and assign its maximum score.

##### E: Batch proportion imbalance

The dataset contains three cell types and two batches. Cell types 1 and 2 have a balanced 500/500 split, while cell type 3 has a 500/100 split. After integration, cell type 1 retains a strong residual batch effect while cell types 2 and 3 show the same small residual batch effect (Fig. 5). An ideal metric should assign a lower score to cell type 1 than to cell types 2 and 3. Importantly, the scores for cell types 2 and 3 should be similar, since their residual batch separation is identical despite different batch proportions.

##### F: Graded batch effect

The dataset contains three cell types and two batches (500 cells per batch per type). After integration, residual batch separation varies across cell types: cell type 1 is well mixed, cell type 2 shows moderate residual batch effect, and cell type 3 shows strong residual batch effect (Fig. 6). An ideal metric should assign clearly separated scores across the three types, with a high score for cell type 1, an intermediate score for cell type 2, and a low score for cell type 3.

##### G: Progressive mixing

The dataset contains two cell types and two batches. Cell type 1 has a strong residual batch effect that remains fixed throughout the scenario. For cell type 2, which starts with a mild residual batch effect, we simulate progressive improvement in batch mixing by gradually shifting the batch 2 cluster toward batch 1 in five equal steps, corresponding to 0%, 25%, 50%, 75%, and 100% of the total distance between the cluster centers (Supplementary Fig. S2). An ideal metric should respond monotonically to this progression for cell type 2, assigning increasing scores as the clusters become more overlapping, while maintaining consistently low scores for cell type 1.

#### Non-Globular Batch Geometries (H–I)

These two scenarios examine whether metrics are sensitive to elliptical batch cluster shapes, covering both separated and intersecting configurations.

##### H: Separated elliptical batches

The dataset contains one cell type and two batches, each forming an elongated elliptical cluster. Two cases are considered: in the first, the two ellipses are close to each other but do not overlap (Supplementary Fig. S4a); in the second, they are distant (Supplementary Fig. S4b). Both cases represent poor integration, but the distant case is more severe. An ideal metric should report low scores in both cases and assign a lower score to the distant case than to the close case. *I: Intersecting elliptical batches.* The dataset again contains one cell type and two batches forming elongated elliptical clusters, but in this scenario the ellipses are oriented at an angle and partially intersect. Two cases are considered: low overlap (Supplementary Fig. S4c) and high overlap (Supplementary Fig. S4d). Because higher overlap indicates weaker residual batch separation, an ideal metric should assign a higher score to the high-overlap case than to the low-overlap case.

#### Concentric and Off-Center Geometries (J–L)

These three scenarios examine metric behavior when batches form concentric or off-center ring-like structures, which arise in practice when one batch forms a subpopulation of another in the embedding space.

##### J: Concentric batches

The dataset contains one cell type and two batches. The outer batch forms a fixed ring with a standard deviation of 2.00, while the inner batch is concentric and its spread is varied across three levels: 0.20 (Supplementary Fig. S5a), 1.00 (Supplementary Fig. S5d), and 2.00 (Supplementary Fig. S5g). At low spread, the inner batch is tightly concentrated and well separated from the outer ring, representing poor mixing. As the spread increases, the inner batch expands to fill the ring, and the two batches become increasingly overlapping. An ideal metric should assign increasing scores across the three spread values, approaching its maximum at spread 2.00.

##### K: Off-center inner batch

This scenario is identical to J in structure but the center of the inner batch is offset from the center of the outer ring (Supplementary Fig. S5b, e, h for spreads 0.20, 1.00, and 2.00, respectively). Because the offset prevents the two batches from fully overlapping even at large spread values, scores should increase with spread but not approach the maximum. An ideal metric should reflect this geometric constraint and plateau below its maximum at the largest spread value.

##### L: Two off-center inner batches

This scenario extends K by replacing the single off-center inner batch with two non-concentric inner batches, each with its own offset center (Supplementary Fig. S5c, f, i for spreads 0.20, 1.00, and 2.00, respectively). The increased separation between the inner and outer batches means that scores should be lower than in K across all spread values. Ideally, the metric should assign scores that are consistently lower than the corresponding scores in K, correctly reflecting the greater degree of residual batch separation.

### 5.2 Real datasets

We evaluated on four real single-cell datasets (Table 3), used in two settings: bench-marking integration methods on two atlas-level datasets from [15], and evaluating the metrics across successive integration stages on the BMMC [29] and HBCA [30] datasets.

**Table 3:** Real datasets used in this work.

| Dataset | Cells | Batches | Cell types | Integration stages |
| --- | --- | --- | --- | --- |
| Human pancreas | 16,382 | 9 | 14 | N/A |
| Human lung | 32,472 | 16 | 17 | N/A |
| BMMC | 69,249 | 13 | 22 | unintegrated, suboptimal, effective, optimized |
| HBCA | 51,367 | 16 | 10 | unintegrated, suboptimal, effective |

#### 5.2.1 Integration method benchmarking

We benchmarked seven integration methods on the human pancreas and human lung atlas datasets of [15], assembled from five and two public RNA datasets respectively. The methods are: a Python implementation of MNN [6] that supports parallel execution across multiple CPUs,^1^ Scanorama [12], ComBat [4], scANVI [28], scVI [9], Harmony [7], and scGen [14].

#### 5.2.2 Metric evaluation across integration stages

We further evaluated the metrics on two multimodal atlas-level datasets: Bone Marrow Mononuclear Cells (BMMC) [29], prepared for the NeurIPS 2021 challenge, and the Human Breast Cell Atlas (HBCA) [30]. For both, the RNA modality was extracted and integrated with the liam method [31] to produce a series of integration settings of increasing batch removal, which were used to evaluate the BRAS metric [24]. BMMC carries a nested batch structure over donor and data-generation site, and was integrated under three settings (suboptimal, effective, optimized); HBCA was integrated under two (suboptimal, effective). We use these, together with the unintegrated baseline, to test the metrics across four integration stages for BMMC and three for HBCA.

### 5.3 Preprocessing

Before computing metric scores, every dataset was reduced to a 50-dimensional PCA embedding. For the real datasets, we followed the scIB preprocessing pipeline [15]: we selected the top 2000 highly variable genes (HVGs) with Scanpy [33] (Cell Ranger flavor) and then performed PCA [34]. The simulated datasets contain no genelevel counts, so HVG selection does not apply; we instead applied PCA directly to their 2000-dimensional projected observations. Unless stated otherwise, metrics were computed on the first 50 principal components.

### 5.4 Evaluation Metrics

Throughout this section, cells are indexed by *i* and *j*. Each cell *i* has an associated cell type *c_i_* and batch label *b_i_*, and is represented by an embedding vector *x_i_* (e.g., the first 50 principal components). We denote by *N_i_* the *k*-nearest neighbors of cell *i* and by I the set of all cells. Let *C_c_* denote the set of all cells of type *c*, *C_c,b_* the set of cells of type *c* in batch *b*, *B_c_* the set of batches containing cell type *c*, and *B* the total number of batches. We compare several widely used batch integration evaluation metrics described below. We report metric scores at two levels of granularity. For the simulated scenarios (where each cell type is constructed to exhibit a different level of residual batch effect, so that per-cell-type scores are informative), we report the percell-type score of each metric that is defined per cell type. For the real datasets, we report a single dataset-level score per metric, aggregated according to each metric’s original design: iLISI, kBET, GC, and ASW take the mean over all cells; CiLISI takes a size-weighted mean across cell types; sBEE macro-averages across cell types (Section 5.5); and PCR is computed globally.

#### Local Inverse Simpson Index (iLISI)

iLISI [7] measures batch diversity in the neighborhood of each cell. For a cell *i* with kernel-weighted local batch probabilities *p_b_*(*i*), the score is

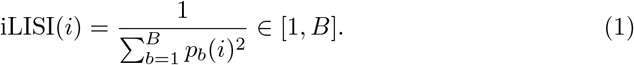

Higher values indicate more balanced batch representation, while values near 1 indicate dominance of a single batch. We rescale to [0, 1] as

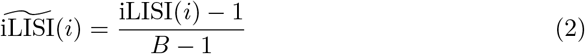

and report the mean 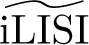 per cell type. We use the implementation of [7] with perplexity 30.

##### Cell Type Integration Local Inverse Simpson Index (CiLISI)

CiLISI [23, 24] is a celltype-adjusted version of iLISI. The rescaled iLISI score 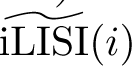 is computed per cell, then averaged within each cell type, and finally summarized as a weighted mean across cell types weighted by cell type size, yielding a single scalar for the entire dataset.

##### Average Silhouette Wi4dth (ASW)

ASW [20], adapted to batch correction by [19] and refined by [15], measures how well batch labels are mixed within biologically defined cell-type clusters. For each cell *i*, the silhouette score with respect to batch labels is

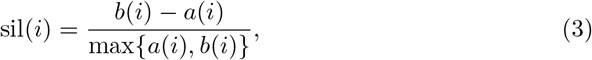

where *a*(*i*) is the mean distance to cells of the same batch within the same cell type, and *b*(*i*) is the mean distance to cells of the nearest neighboring batch within the same cell type. Following [15], we transform to

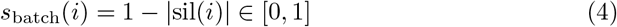

so that higher values indicate better mixing, and report the mean over all cells of each type. ASW requires at least two batches per cell type and is set to NaN otherwise.

##### Batch Removal Adapted Silhouette (BRAS)

BRAS [24] is a modified version of batch ASW. While [15] ASW defines *b*(*i*) as the mean distance to the nearest neighboring batch (within the same cell type), BRAS redefines *b*(*i*) as the mean distance to all cells from other batches within the same cell type, making it robust to the nearest-cluster issue. BRAS also uses cosine rather than Euclidean distance for higher discriminative power.

##### k-Nearest Neighbor Batch Effect Test (kBET)

kBET [19] evaluates whether the local batch composition around each cell matches the global batch proportions using a Pearson *χ*^2^ goodness-of-fit test. Let *p_b_* denote the global proportion of batch *b*. For cell *i*, the statistic is

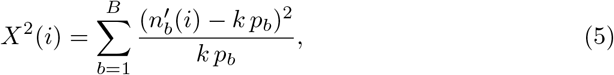

where 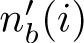 is the number of neighbors from batch *b*. We report the acceptance rate per cell type as

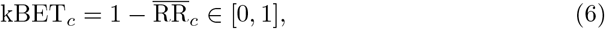

where RR*_c_*is the mean rejection rate over *n*_repeat_ = 100 subsampled trials (approximately 10% of cells) with significance threshold *α* = 0.05.

##### Graph Connectivity (GC)

GC [15] measures the fraction of cells of each type that belong to the largest connected component (LCC) of the *k*-nearest neighbor graph after integration:

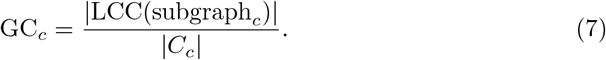

Values close to 1 indicate that cells of the same type form a single connected component.

##### Principal Component Regression (PCR)

PCR [15, 19] quantifies how much of the total variance is associated with batch across principal components. Let **Z** ∈ {0, 1}*^n×B^* be the one-hot batch design matrix for the *n* cells, and let **f***_ℓ_* ∈ R*^n^* collect the scores of all cells on principal component PC*_ℓ_*. For each component we fit the linear regression

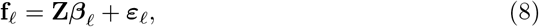

and measure the fraction of that component’s variance explained by batch through the coefficient of determination

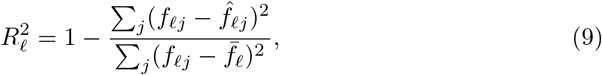

where *f̂_ℓj_* are the fitted values and *f̄_ℓ_* is the mean score. Weighting each component by the proportion of total variance it carries, 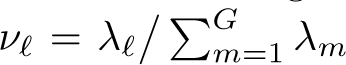 (with *λ_ℓ_* the variance of PC*_ℓ_*), the batch-associated variance is

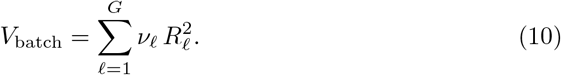

We report 1 − *V*_batch_ ∈ [0, 1], where higher values indicate better batch mixing. Unlike the other metrics, PCR is computed globally across all cells rather than per cell type. We apply the scIB implementation to the shared 50-dimensional PCA embedding, so the sum runs over the *G* = 50 principal components.

### 5.5 The Proposed Metric: sBEE

We introduce sBEE (Fig. 1), a per-cell metric that evaluates batch mixing within each cell type from two complementary views: local neighborhood composition and cross-batch distances. Scores lie in [0, 1], with higher values indicating better integration.

To capture local batch mixing, we compare the batch composition of the *k*-nearest neighbors of cell *i* with the global batch distribution for cell type *c_i_*:

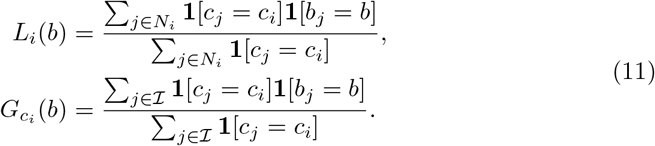

We quantify the discrepancy between these two distributions, *L_i_*and *G_c__i_*, using the Jensen–Shannon distance [35], referred to below as *D*_JS_. JS distance is bounded in [0, 1]; smaller values indicate better agreement between local and global batch composition. For each cell type *c*, the effective neighborhood size is defined as *k_c_* = min(*k,* |*C_c_*| − 1), where |*C_c_*| is the number of cells of type *c*. Thus, the neighborhood of a cell never extends beyond the available cells of the same type, and the effective neighborhood size is adaptive across cell types.

For each cell *i*, we define two same-type reference sets

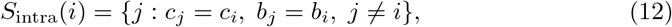

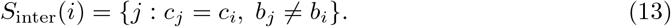

Using the embedding *x_i_*, the median intra-and inter-batch distances are

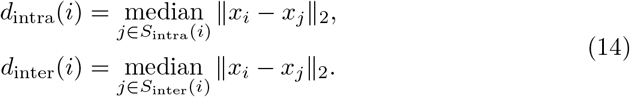

We use the median rather than the mean to reduce sensitivity to outlier cells. We quantify cell-type batch separation by comparing distances from a cell to same-type cells within the same batch and to those from different batches:

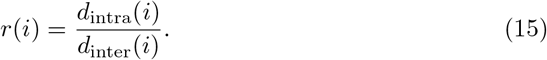

Values *r*(*i*) ≈ 1 indicate well-mixed batches, whereas *r*(*i*) ≪ 1 indicates residual batch separation. To prevent over-penalization when intra-batch distances exceed inter-batch distances, we cap *r*(*i*) = min(*r*(*i*), 1). Cells with |*S*_intra_(*i*)| = 0 are excluded (*r*(*i*) = NaN), and when a cell type appears in only one batch we set *r*(*i*) = 1.

The neighborhood mixing component and the batch-separation component (based on the ratio *r*(*i*) defined above) are, respectively,

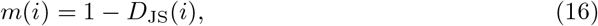

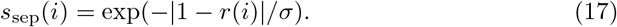

The parameter *σ >* 0 controls the sensitivity of the separation component to deviations of *r*(*i*) from the ideal value of 1; based on a sensitivity analysis across scenarios (Supplementary Fig. S8), we set *σ* = 0.15, the value that preserves score orderings while maintaining discrimination among configurations with differing degrees of separation.

The final per-cell score is obtained using the harmonic mean

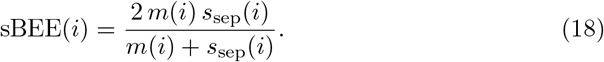

Cell-type scores are obtained by macro-averaging cell-level sBEE scores across batches within each cell type, ensuring equal batch contribution at the cell-type level. The overall dataset score is then obtained by macro-averaging across all cell-type scores, ensuring each cell type contributes equally regardless of its size.

## Acknowledgements

This work is supported by the Scientific and Technological Research Council of Türkiye (TUÜBIİTAK), under grant numbers 121E491 given to H.K and 125N682 given to O.T.

## Data availability

The sBEE source code is publicly available on GitHub ((https://github.com/tastanlab/sBEE). The simulated evaluation scenarios are available on Zenodo (https://doi.org/10.5281/zenodo.21031593) [36]. The pancreas datasets are available on Zenodo (https://doi.org/10.5281/zenodo.21069083) [37]. The lung datasets are available on Zenodo (https://doi.org/10.5281/zenodo.21070566) [38]. The embeddings of the BMMC and HBCA datasets are available on Zenodo (https://doi.org/10.5281/zenodo.15705865) [39].

## Authors’ contributions

O. T. and H. K. conceived the study, supervised the project, and contributed to writing the manuscript. M. M. generated the data, implemented the method, performed the analyses, and contributed to writing the manuscript. A. H. assisted with data generation and analysis. All authors read and approved the final version of the manuscript.

## Ethics approval and consent to participate

Not applicable.

## Competing interests

The authors declare no competing interests.

## Footnotes

1 https://github.com/chriscainx/mnnpy

## Notes

https://github.com/tastanlab/sBEE

### Competing Interest Statement

The authors have declared no competing interest.

### Summary of Updates

Sections were rewritten, figures were remade.

