## Supplementary File for "A Systematic Evaluation of Single-Cell Batch Integration Metrics and sBEE, A New Metric"

### Supplementary Information for "A Systematic Evaluation of Single-Cell Batch Integration Metrics and sBEE, A New Metric"

#### 1 Supplementary Figures

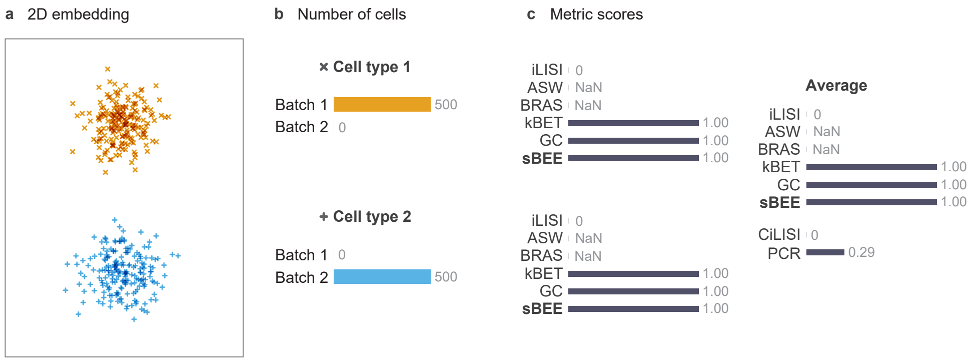


**Fig. S1** a) Simulated 2D embedding for Scenario D (batch-specific cell types). b) Per-cell-type batch composition, where colors represent batches. c) Metric scores. Each cell type has a subplot showing iLISI, ASW, BRAS, kBET, GC, and sBEE, followed by a subplot of their average across cell types together with the dataset-level metrics CiLISI and PCR. CiLISI and PCR (set apart by whitespace) report a single value per scenario rather than per-cell-type scores. Scores range from 0 to 1; higher values indicate better integration.

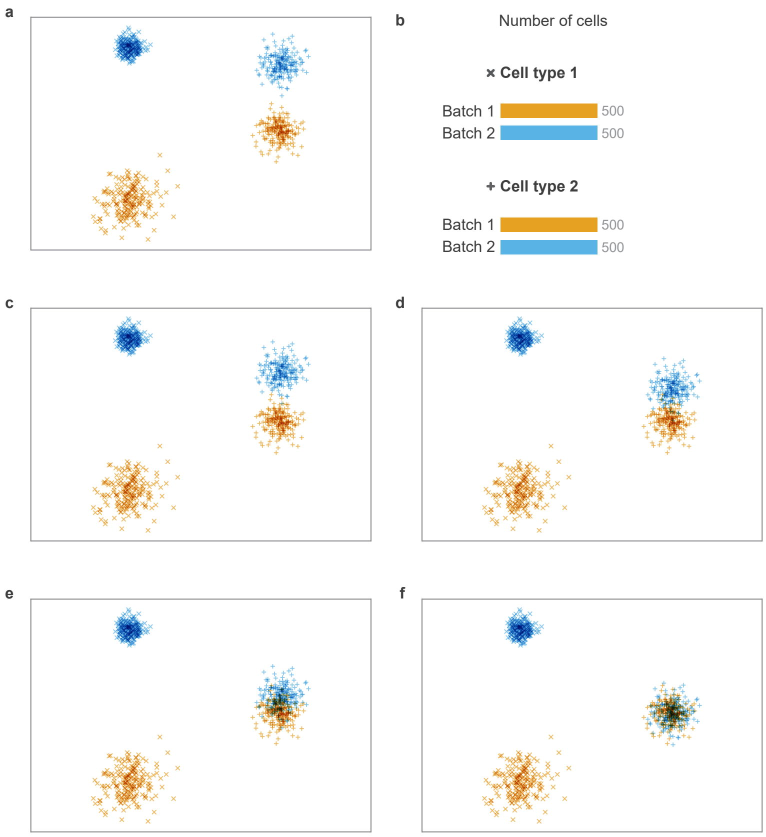


**Fig. S2** (a, c–f) Simulated 2D embedding for Scenario G (progressive mixing). (b) Per-cell-type batch composition, where colors represent batches. Two cell types, two batches; cell type 1 retains strong residual batch separation throughout, while cell type 2 is progressively shifted toward full mixing in five equal steps (0%, 25%, 50%, 75%, and 100% of the total distance between cluster centers).

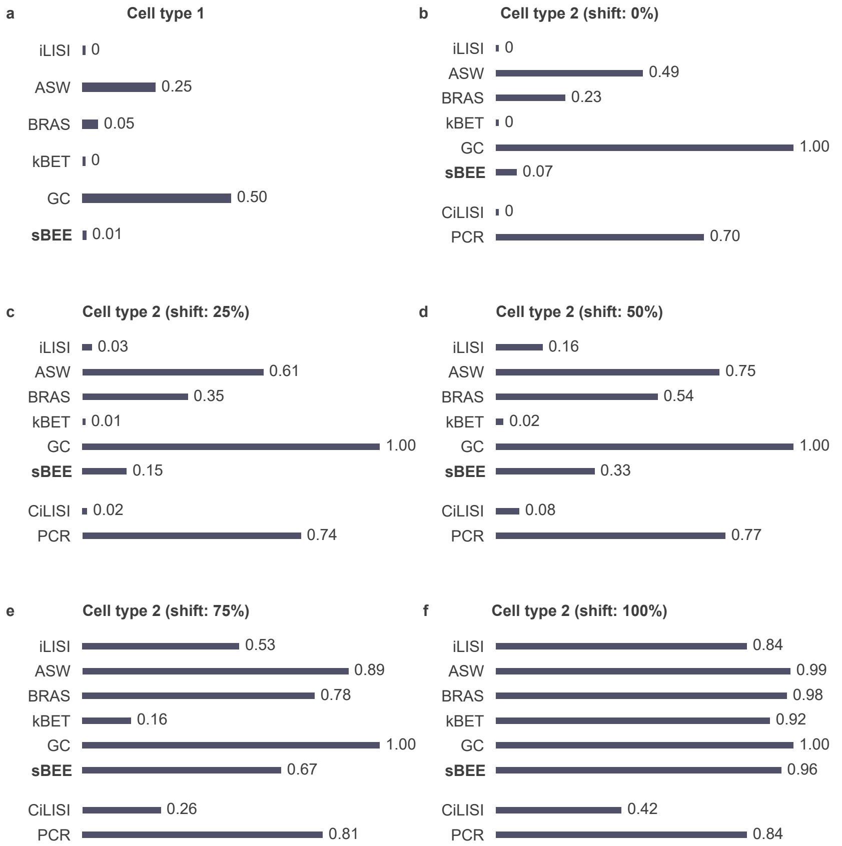


**Fig. S3** Scenario G: Progressive mixing scores. Metric scores across the five progressive mixing steps for cell types 1 and 2. CiLISI and PCR yield a single dataset-level value rather than per-cell-type scores. Scores range from 0 to 1; higher values indicate better integration.

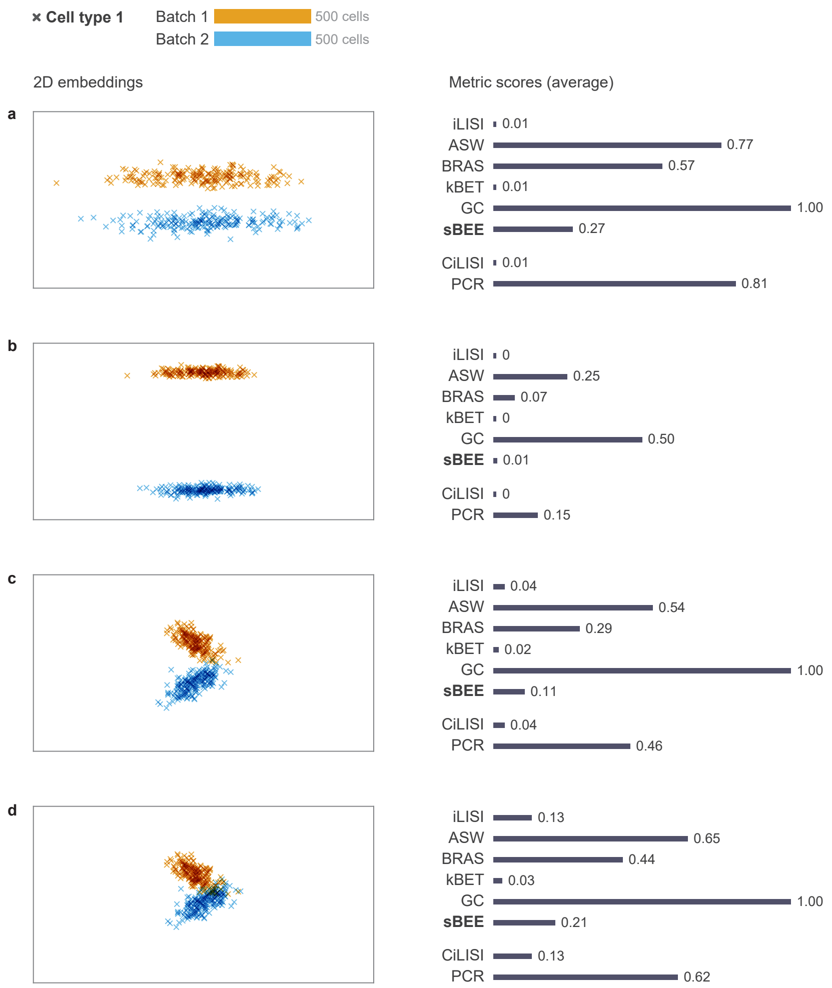


**Fig. S4** Simulated 2D embeddings and corresponding metric scores for Scenarios H and I (non-globular batch geometries). Panels **a** and **b** correspond to Scenario H, and panels **c** and **d** correspond to Scenario I. CiLISI and PCR are set apart by whitespace as dataset-level metrics. Scores range from 0 to 1, higher values indicate better integration.

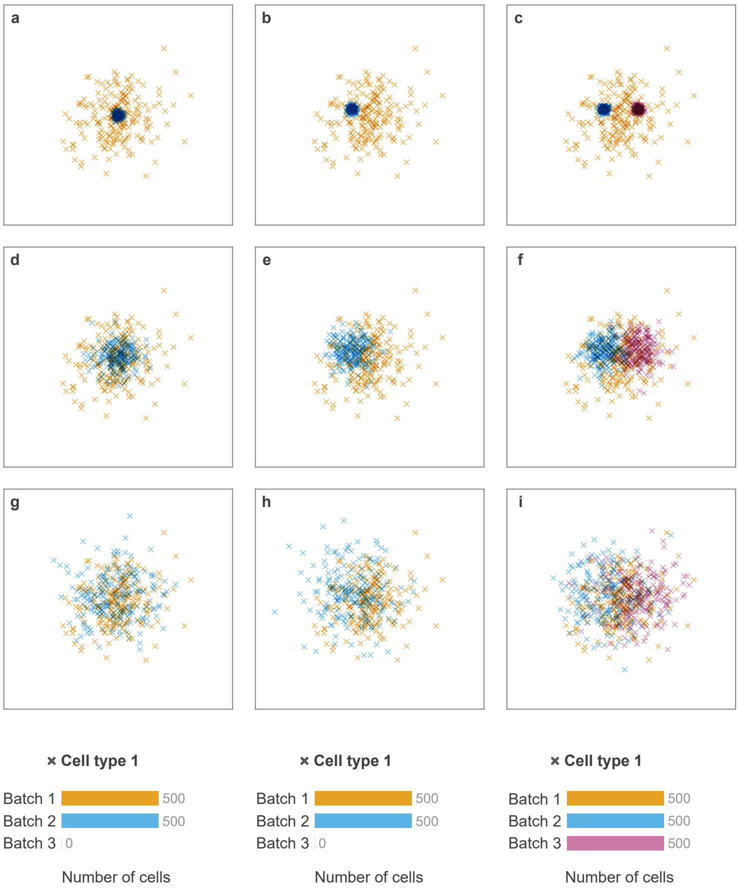


**Fig. S5** Simulated 2D embeddings for three spread values (0.20, 1.00, 2.00) across three geometry types. (a, d, g) Concentric batches (scenario J). (b, e, h) Single off-center inner batch (scenario K). (c, f, i) Two off-center inner batches (scenario L).

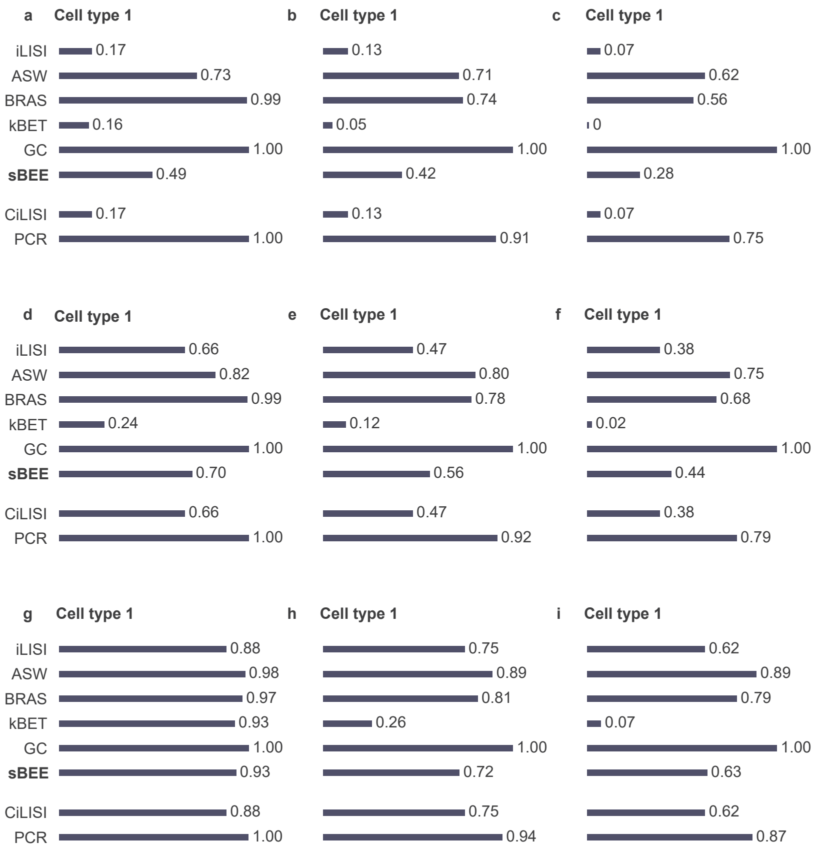


**Fig. S6** Metric scores for concentric and off-center geometries. (a, d, g) Scores for scenario J across three spread values. (b, e, h) Scores for scenario K. (c, f, i) Scores for scenario L. CiLISI and PCR are set apart by whitespace as dataset-level metrics. Scores range from 0 to 1; higher values indicate better integration.

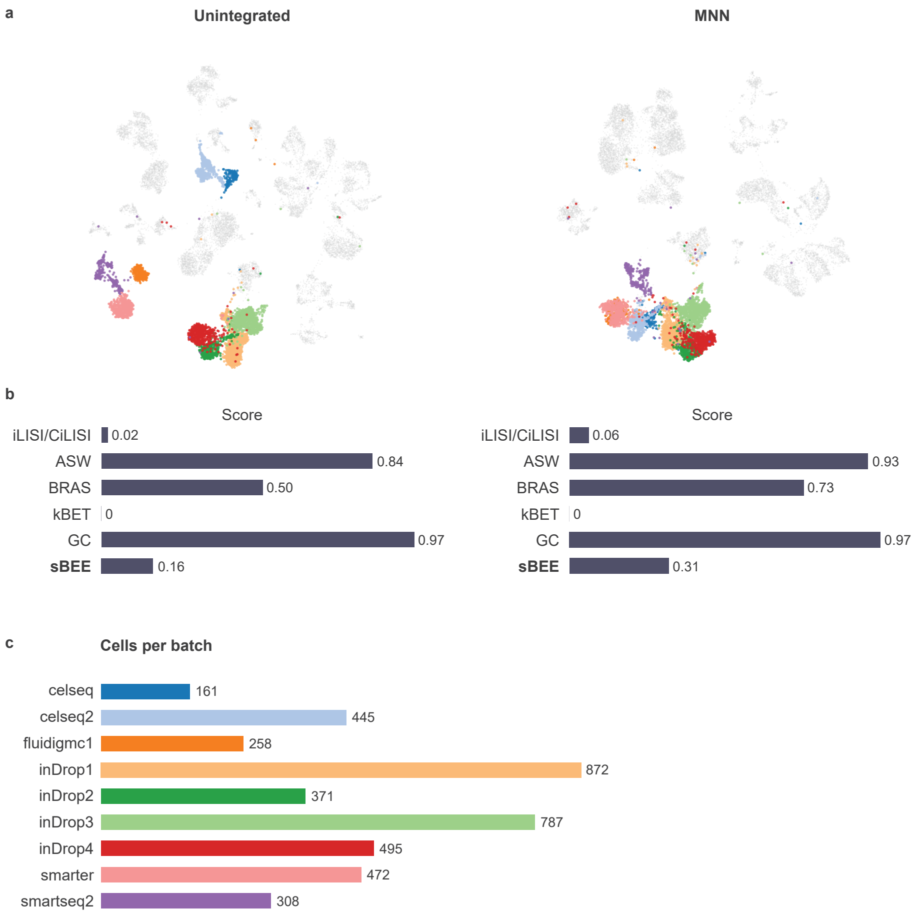


**Fig. S7** Batch mixing of beta cells before and after MNN integration. a) UMAP embeddings of the unintegrated and MNN-integrated pancreas datasets. Beta cells are colored by batch, while all other cell types are shown in grey. b) Batch-correction metrics computed for beta cells. c) Number of beta cells contributed by each batch.

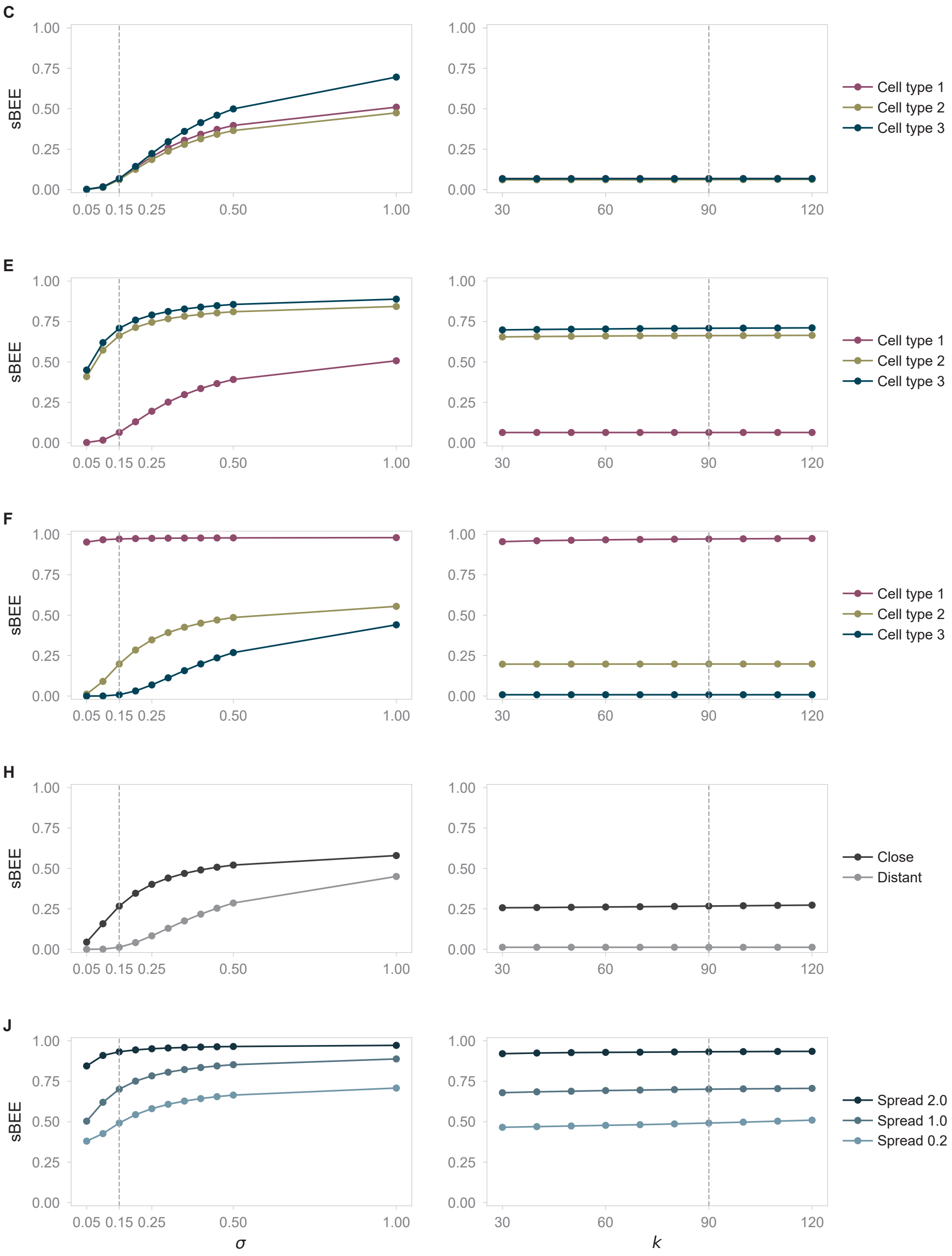


**Fig. S8** sBEE is robust to its parameter choices. For five representative scenarios (C, E, F, H, J), sBEE cell-type scores are shown as a function of the separation-component scale $\sigma$ (left, with $k = 90$) and the neighborhood size $k$ (right, with $\sigma = 0.15$). Dashed lines mark the default values. Scores are essentially invariant to $k$ across the range 30–120, while $\sigma$ rescales the scores without changing the ordering or pass/fail verdict over a broad range; the default $\sigma = 0.15$ keeps correctly-low scores low while preserving discrimination among more separated configurations.

### 2 Supplementary Tables

**Table 1** Simulation scenarios. Data are drawn from a 2D Gaussian; each (cell type, batch) pair has a distinct mean vector $\boldsymbol{\mu}_{c,b} \in \mathbb{R}^2$ and standard deviation $\sigma_{c,b}$.

| Scenario | Mean vectors $\boldsymbol{\mu}_{c,b}$ | Std. devs. $\sigma_{c,b}$ |
|---|---|---|
| A | $\boldsymbol{\mu}_{1,*} = [1, 10]$, $\boldsymbol{\mu}_{2,*} = [5, 1]$ | $\sigma_{*,*} = 0.6$ |
| B | $\boldsymbol{\mu}_{1,1} = [1, 4]$, $\boldsymbol{\mu}_{1,2} = [1, 3.5]$, $\boldsymbol{\mu}_{1,3} = [5, 3]$ | $\sigma_{*,*} = 0.6$ |
| C | $\boldsymbol{\mu}_{1,1} = [1, 5]$, $\boldsymbol{\mu}_{1,2} = [1, 1]$,<br>$\boldsymbol{\mu}_{2,1} = [10, 5]$, $\boldsymbol{\mu}_{2,2} = [10, 1]$,<br>$\boldsymbol{\mu}_{3,1} = [20, 5]$, $\boldsymbol{\mu}_{3,2} = [20, 1]$ | $\sigma_{*,*} = 0.6$ |
| D | $\boldsymbol{\mu}_{1,1} = [1, 1]$, $\boldsymbol{\mu}_{2,2} = [5, 5]$ | $\sigma_{*,*} = 0.6$ |
| E | $\boldsymbol{\mu}_{1,1} = [1, 5]$, $\boldsymbol{\mu}_{1,2} = [1, 1]$,<br>$\boldsymbol{\mu}_{2,1} = [10, 4]$, $\boldsymbol{\mu}_{2,2} = [10, 3]$,<br>$\boldsymbol{\mu}_{3,1} = [20, 4]$, $\boldsymbol{\mu}_{3,2} = [20, 3]$ | $\sigma_{*,*} = 0.6$ |
| F | $\boldsymbol{\mu}_{1,*} = [10, 10]$,<br>$\boldsymbol{\mu}_{2,1} = [15, 9]$, $\boldsymbol{\mu}_{2,2} = [15, 11]$,<br>$\boldsymbol{\mu}_{3,1} = [20, 5]$, $\boldsymbol{\mu}_{3,2} = [20, 15]$ | $\sigma_{*,*} = 0.3$ |
| G | $\boldsymbol{\mu}_{1,1} = [1, 1]$, $\boldsymbol{\mu}_{1,2} = [1, 10]$,<br>$\boldsymbol{\mu}_{2,1} = [10, 5]$, $\boldsymbol{\mu}_{2,2} = [10, 9]$ | $\sigma_{1,1} = 0.9$, $\sigma_{1,2} = 0.4$,<br>$\sigma_{2,1} = 0.6$, $\sigma_{2,2} = 0.6$ |
| H | $\boldsymbol{\mu}_{1,1} = [0, 2]$, $\boldsymbol{\mu}_{1,2} = [0, -2]$ (lower sep.)<br>$\boldsymbol{\mu}_{1,1} = [0, 10]$, $\boldsymbol{\mu}_{1,2} = [0, -10]$ (higher sep.) | $\sigma_{*,*} = 0.5^{\dagger}$ |
| I | $\boldsymbol{\mu}_{1,1} = [-2, 1]$, $\boldsymbol{\mu}_{1,2} = [-2, -1]$ (lower overlap)<br>$\boldsymbol{\mu}_{1,1} = [-2, 1]$, $\boldsymbol{\mu}_{1,2} = [-1, 0]$ (higher overlap) | $\sigma_{*,*} = 1^{\dagger}$ |
| J | $\boldsymbol{\mu}_{*,*} = [0, 0]$ | $\sigma_{1,1} = 2.0$,<br>$\sigma_{1,2} = \{0.2,\ 1.0,\ 2.0\}$ |
| K | $\boldsymbol{\mu}_{1,1} = [0, 0]$, $\boldsymbol{\mu}_{1,2} = [-1.5, 0.5]$ | $\sigma_{1,1} = 2.0$,<br>$\sigma_{1,2} = \{0.2,\ 1.0,\ 2.0\}$ |
| L | $\boldsymbol{\mu}_{1,1} = [0, 0]$<br>$\boldsymbol{\mu}_{1,2} = [-1.5, 0.5]$<br>$\boldsymbol{\mu}_{1,3} = [1.5, 0.5]$ | $\sigma_{1,1} = 2.0$,<br>$\sigma_{1,2} = \{0.2,\ 1.0,\ 2.0\}$,<br>$\sigma_{1,3} = \{0.2,\ 1.0,\ 2.0\}$ |

$^{\dagger}$ Clusters are elliptical: points are first sampled from an isotropic Gaussian with the given $\sigma$, then the $x$-coordinate is scaled by a factor of 8.
